# Proximity-based labeling reveals DNA damage-induced N-terminal phosphorylation of fused in sarcoma (FUS) leads to distinct changes in the FUS protein interactome

**DOI:** 10.1101/2021.06.11.448082

**Authors:** Michelle A. Johnson, Thomas A. Nuckols, Paola Merino, Pritha Bagchi, Srijita Nandy, Jessica Root, Georgia Taylor, Nicholas T. Seyfried, Thomas Kukar

**Affiliations:** Department of Pharmacology and Chemical Biology, Emory University, School of Medicine, Atlanta, GA; Center for Neurodegenerative Disease, Emory University, School of Medicine, Atlanta, GA; Emory Integrated Proteomics Core, Emory University, School of Medicine, Atlanta, GA; Department of Neurology, Emory University, School of Medicine, Atlanta, GA; Department of Biochemistry, Emory University, School of Medicine, Atlanta, GA

**Keywords:** Fused in sarcoma (FUS), phosphorylation, proteomics, translation, protein-protein interaction, protein translocation, amyotrophic lateral sclerosis (ALS) (Lou Gehrig disease

## Abstract

Cytoplasmic accumulation of the RNA/DNA binding protein, fused in sarcoma (FUS), into inclusions is a common hallmark of frontotemporal lobar degeneration (FTLD) and amyotrophic lateral sclerosis (ALS) pathology. We have previously shown that DNA damage can trigger the cytoplasmic accumulation of an N-terminally phosphorylated FUS. However, the functional consequences of N-terminal FUS phosphorylation are unknown. To gain insight into this question, we utilized proximity-dependent biotin labeling via ascorbate peroxidase 2 (APEX2) paired with mass-spectrometry (MS) to investigate whether N-terminal phosphorylation shifts the FUS protein-protein interaction network (interactome), and subsequently, its function. We report the first comparative analysis of the interactomes for three FUS variants: homeostatic wild-type FUS (FUS WT), a phosphomimetic variant of FUS (a proxy for N-terminally phosphorylated FUS, FUS PM), and a toxic FUS P525L mutant (a mutation that causes juvenile ALS, FUS P525L). Data are available via ProteomeXchange with identifier PXD026578. We demonstrate that compared to FUS WT and FUS P525L, the FUS PM interactome uniquely enriches for a set of cytoplasmic proteins that mediate mRNA metabolism and translation and nuclear proteins involved in spliceosome and DNA repair functions, respectively. We further identify and validate three proteins, VPS35, MOV10, and CLTA, as novel interacting partners of all three FUS variants. Lastly, we provide functional evidence that N-terminally phosphorylated FUS may disrupt homeostatic translation and steady state levels of specific mRNA transcripts. Taken together, these results highlight phosphorylation as a unique modulator of the FUS interactome and function.

## Introduction

Frontotemporal lobar degeneration (FTLD) is a neurodegenerative disease characterized by atrophy of the frontal and temporal lobes. The clinical manifestation of FTLD is frontotemporal dementia (FTD) (1). FTD is a heterogenous group of clinical disorders that either results in alterations to behavior and personality or impairments in language comprehension and communication (1, 2). Pathological and genetic similarities between FTD and another neurodegenerative disease, amyotrophic lateral sclerosis (ALS), suggest that FTD and ALS exist on a disease spectrum (3–6). ALS is a progressive motor neuron disease characterized by degeneration of upper and lower motor neurons (3, 7). While ALS typically targets a different neuronal population compared to FTLD, pathology in a subset of both diseases is linked to the abnormal aggregation of the fused in sarcoma (FUS) protein (8–11).

FUS is a pleiotropic RNA/DNA binding protein involved in gene transcription, DNA-repair pathways, mRNA splicing, mRNA transport, and stress granule assembly (9,12–20). In FTLD-FUS and ALS-FUS postmortem tissue, FUS is found in cytoplasmic inclusions or aggregates in neurons and glia (21–24). This cytoplasmic accumulation is directly correlated with toxicity (25, 26). Cellular dysfunction related to FUS aggregation is thought to be driven by novel gain-of-functions that trigger cellular death (25–30). As such, understanding how these gain-of-functions induced by cytoplasmic FUS contribute to toxicity will be necessary to understanding disease pathogenesis and developing targeted therapies.

Neuronal cytoplasmic inclusions that contain FUS occur in ∼10% of FTD cases and ∼5% of ALS cases (1,31,32). Genetic mutations in *FUS* typically cause ALS and are rarely associated with FTLD (21,24,33,34). Thus, the proximal cause of FUS pathology in FTLD is unknown. One possibility is that FUS pathology is caused by exposure to an environmental toxin or dysregulated post-translational modifications (PTMs), such as phosphorylation or methylation (15,35–41).

Phosphorylation is a reversible PTM that regulates the function of numerous proteins in the cell (42). Abnormal or dysregulated protein phosphorylation is a common feature of many neurodegenerative disorders, including FTLD and ALS (43, 44). FUS can be phosphorylated at multiple N- and C-terminal residues, but the functional consequence of these modifications remain largely unexplored (36,45–48). Our lab discovered that phosphorylation of 12 specific N-terminal residues in FUS by the DNA-dependent protein kinase (DNA-PK) causes the cytoplasmic accumulation of phosphorylated FUS (40,47,49). This cascade is triggered by double strand DNA breaks (DSB). Various studies have found that FTLD and ALS exhibit markers of DNA damage. Given this, the cytoplasmic re-localization of FUS induced by N-terminal phosphorylation may contribute to pathology in a subset of FTLD and ALS cases (38,47,50,51). However, it remains unclear whether N-terminal phosphorylation alters FUS function. Alterations in FUS function are commonly studied at the level of the proteome and interactome (14,52–54). Even so, no study has directly mapped how a PTM may shift the interactome of FUS. Therefore, we aimed to elucidate how the FUS protein interactome changed in response to phosphorylation at these 12 key N-terminal residues.

We performed proximity-mediated biotin labeling coupled with label-free mass spectrometry (MS) to determine whether N-terminal phosphorylation alters the protein binding partners of FUS (55). Chemically induced DSBs lead to robust phosphorylation of FUS but are toxic to cells making proteomic analysis challenging (Deng *et al.*, 2014b). To overcome this hurdle, we focused our analysis on a phosphomimetic variant of FUS (FUS PM) that mimics the cytoplasmic localization caused by DSBs (47, 49). We engineered synthetic genes that fused APEX2 to human wild-type FUS (FUS WT), FUS PM, or the ALS-linked mutant P525L (FUS P525L) to enable proximity-dependent biotinylation of potential protein binding partners (55). Differential expression analysis revealed that the FUS PM interactome was enriched for cytoplasmic proteins involved in “mRNA catabolic process”, “translation initiation”, and “stress granule assembly” over FUS WT. In contrast, the FUS PM interactome was enriched for nuclear proteins involved in functions such as “spliceosome”, “ribonucleoprotein complex biogenesis”, and “covalent chromatin modification” compared to FUS P525L. We found that cells expressing FUS PM exhibited functional alterations in the steady-state levels of certain mRNAs and global translation. Taken together, these data suggest that phosphorylation results in a novel FUS interactome that exists between the pathogenic FUS P525L ALS-linked mutation, and the homeostatic functions of FUS WT. Our analysis is the first comprehensive study of how a disease-relevant post-translational modification in FUS may shift its protein interactome towards a disease state. Findings from these studies will inform how phosphorylation of FUS and an ALS-linked FUS mutation contribute to neurodegeneration.

## Results

### APEX2 tagged Phosphomimetic FUS (FUS PM) recapitulates p-FUS localization phenotype

FUS dysfunction is involved in FTD and ALS disease pathogenesis. However, many fundamental aspects of the function and regulation of FUS are unknown. For example, it remains unclear how phosphorylation of FUS, or the presence of ALS-associated mutations, alters the function of FUS and associated pathways. To gain insight into these questions, we set out to define the protein binding network, or interactome, of FUS by performing proximity labeling mediated by ascorbate peroxidase 2 (APEX2). We genetically fused APEX2 to the N-terminus of three FUS protein variants via a (GGGS)^3^ linker to generate three Twin-Strep-tagged® constructs: 1) wild-type human FUS (FUS WT), 2) phosphomimetic FUS (FUS PM), and 3) the ALS-linked P525L mutant FUS (FUS P525L) (Figure 1A). FUS PM was generated by substituting the 12 consensus S/T_Q residues, which are phosphorylated by DNA-PK following DSB, with the negatively charged amino acid aspartate (47, 49). The FUS P525L mutation was first identified in 2012 and causes a severe form of juvenile ALS (56, 57). FUS P525L robustly increases cytoplasmic localization of FUS and alters the transcriptome, proteome, and the spliceosome in multiple model systems (12,58,59). Therefore, the APEX2-FUS P525L mutant 1) served as a positive control for FUS cytoplasmic localization, 2) provided insight into the pathogenic nature of ALS-linked mutations, and 3) was a useful comparison to determine if FUS PM resembles a pathogenic phenotype (45, 50).

**Figure 1.**
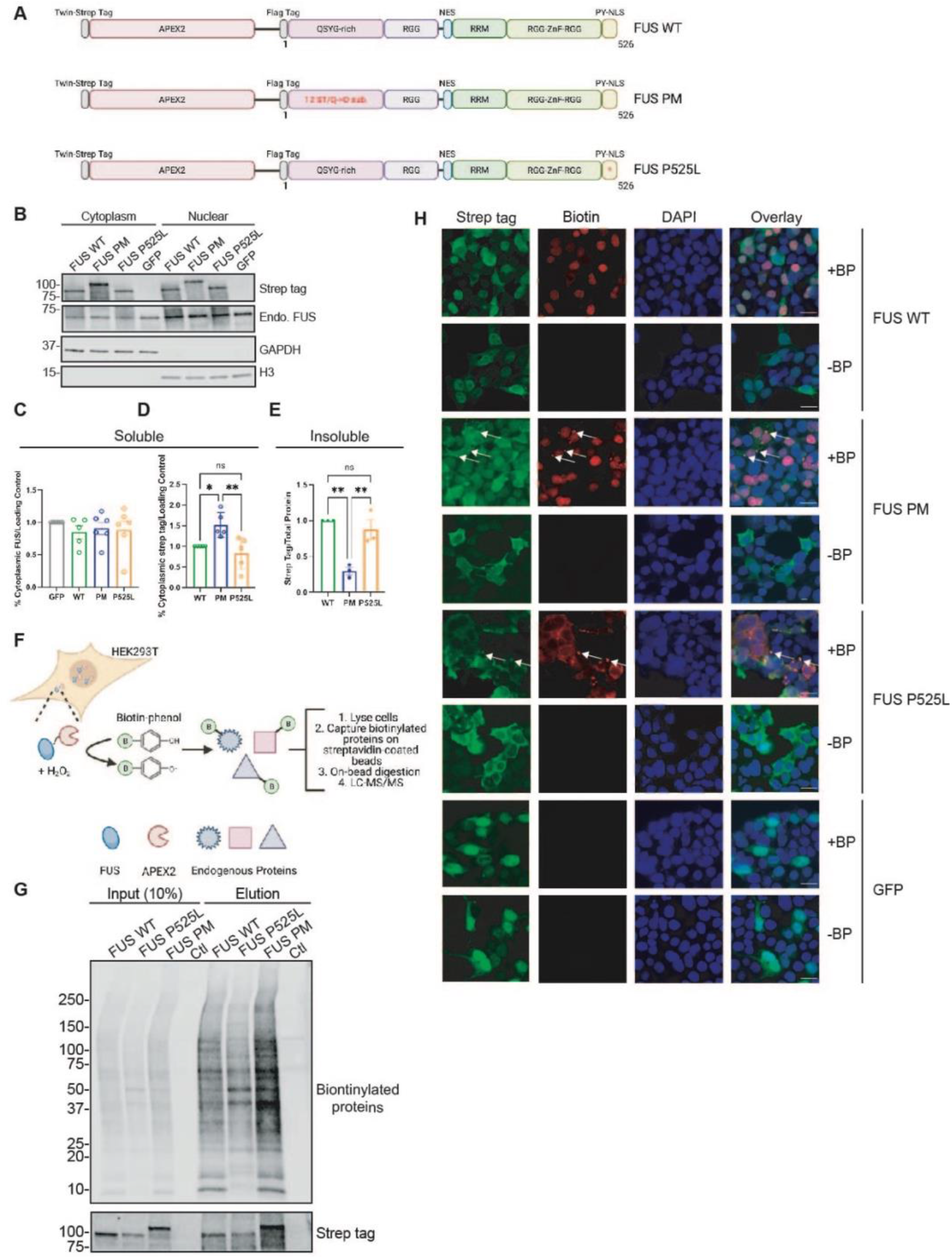
Biotinylation pattern induced by APEX2 is dependent on FUS variant localization and solubilization. (A) Graphical representation of the domain structures within the three APEX2-FUS fusion constructs. Each construct contains a twin-strep tag for identification in downstream applications, APEX2, a linker sequence and a variant of human full-length FUS. The three FUS variants are: wildtype FUS (FUS WT), phosphomimetic FUS where either serine or threonine at the 12 DNA-PK consensus sites (S/T-Q) were mutated to aspartate (D), and pathogenic P525L mutant FUS. Created with BioRender.com. (B) HEK293T cells expressing the three APEX2-FUS fusion constructs were fractionated for cytoplasm and nuclear fractions. GAPDH and H3 were used as markers for cytoplasmic and nuclear fractions, respectively. (C) Quantification of (B) for the percentage of strep-tagged APEX2 fusion proteins found within the soluble cytoplasmic fraction and normalized to loading control. (D) Quantification of (B) for the percentage of endogenous FUS found within the soluble cytoplasmic fraction and normalized to loading control. (E) Quantification of the proportion of strep-tagged APEX2 fusion proteins within the detergent insoluble fraction and normalized to total protein (Immunoblot not shown). (F) Schematic representation of APEX2 proximity labeling and biotin enrichment in presence of H_2_O_2_ and biotin-phenol. Created with BioRender.com. (G) Enrichment of biotinylated proteins from HEK293T cells expressing various APEX2 constructs and treated with biotin-phenol and H_2_O_2_. Input is 1% of sample loaded onto magnetic beads coated with streptavidin; Elute is 10% of sample eluted off beads. Samples are wildtype FUS (FUS WT), P525L FUS (FUS P525L), Phosphomimetic FUS (FUS PM), and non-transfected control (CTL). Input and elution were analyzed for biotinylated proteins (streptavidin) and Twin-Strep-tag® (strep tag). (H) Immunostaining of HEK293T cells expressing the three APEX2-FUS fusion constructs that have been given biotin-phenol (BP) and H_2_O_2_ for twin-strep tag (fusion protein) and streptavidin (biotin). Scale bar represents 20µm.

We first asked if fusion of APEX2 maintained the expected sub-cellular localization of the FUS variants. We expressed the three APEX2 fusion constructs in HEK293T cells and biochemically fractionated cells into a soluble cytoplasmic and nuclear fraction (Figure 1B). Endogenous FUS protein was enriched in the nuclear fraction and the ratio of cytoplasmic/nuclear FUS was unchanged regardless of APEX2-fusion protein expression (Figure 1C). Previously, we reported that the cytoplasmic localization of phosphorylated FUS induced by DSB can be mimicked by phosphomimetic substitution of the 12 consensus DNA-PK phosphorylation sites (S/T_Q) with aspartate (D) (47). As anticipated, a larger proportion of APEX2-FUS PM was found in the cytoplasm compared to the nucleus via western blot (Figure 1D). Western blot analysis also revealed a significant increase in APEX2-FUS WT and APEX2-FUS P525L localized to the soluble nuclear fraction (Figure 1E). FUS-ALS mutations such as P525L typically induce an accumulation of FUS into insoluble cytoplasmic inclusions (22, 32). As such, we examined the insoluble protein fraction and found that APEX2-FUS WT and APEX2-P525L FUS were both significantly increased in the insoluble fraction compared to APEX2-FUS PM. This suggested that a significant fraction of APEX2-FUS WT and APEX2-FUS P252L is detergent insoluble (Figure 1E). Insoluble APEX2-FUS WT or APEX2-P525L protein could localize to either the nucleus or the cytoplasm. Therefore, we next utilized immunofluorescent staining to determine the sub-cellular localization of the APEX2 fusion proteins without relying on detergent-based fractionation. In line with western blot analysis, APEX2-FUS WT was found in the cytoplasm and nucleus. In contrast, both APEX2-FUS PM and APEX2-FUS P525L showed a more pronounced cytoplasmic localization (Figure 1H). Taken together, our data demonstrate that the APEX2 fusion FUS variants localize to expected cellular compartments.

### APEX2-FUS variants exhibit unique biotinylation patterns

To further validate the proximity ligation system, we confirmed that the APEX2 fusion proteins are active and can biotinylate endogenous proteins. APEX2 requires the addition of biotin-phenol and H_2_O_2_ to catalyze the biotinylation of proximal endogenous proteins (Figure 1F). When we treated cells expressing the APEX2-FUS variants with biotin-phenol and H_2_O_2_, we observed robust and variant specific, biotin labeling of endogenous proteins as detected by immunofluorescence with streptavidin (Figure 1H). In contrast, we did not observe biotin labeling in cells that were not treated with biotin-phenol or H_2_O_2_ (Supplementary Figure 1). While APEX2-FUS WT exhibited a mixed nuclear and cytoplasmic localization when immunostained for the Twin-Strep-tag® (Figure 1H), it induced a primarily nuclear biotinylation pattern as determined by colocalization with streptavidin (biotin) and DAPI immunofluorescence. APEX2-FUS PM exhibited a diffuse cytoplasmic localization pattern with biotinylated proteins primarily labeled in the nucleus with interspersed cytoplasmic puncta (white arrows). APEX2-FUS P525L was localized primarily to the cytoplasm and induced biotinylation in the cytoplasm. Negative control cells expressing a strep tagged GFP show no biotinylation following biotin-phenol and H_2_O_2_ addition. These results demonstrate that APEX2-FUS variants exhibit unique and specific patterns of biotinylation.

To identity the variant specific binding partners of APEX2-FUS proteins, we transfected HEK293T cells with three APEX2 constructs (APEX2-FUS WT, APEX2-FUS PM, or APEX2-FUS P525L) for 24 hours. Untransfected HEK293T cells were grown in parallel for 24 hours and served as a control group. All biological groups contained technical replicates done in quadruplicate. We incubated each experimental group of cells with biotin-phenol for 30 minutes followed by H_2_O_2_ for 1 minute to induce biotinylation of proximal endogenous proteins. The reaction was quenched, and lysates were collected (Figure 1G). While control cells did not receive biotin-phenol, they did receive H_2_O_2_ and underwent all downstream processing. Biotinylated proteins were enriched from the cell lysates using streptavidin affinity purification. Western blot analysis of ∼10% of the volume of streptavidin beads confirm enrichment of biotinylated proteins and revealed that each APEX2 FUS variant showed a distinct biotinylation pattern (Figure 1G). The remaining affinity-purified biotinylated proteins were used for unbiased proteomic analysis.

### APEX2-induced biotinylation identities novel binding partners of FUS variants

To identify novel FUS interacting proteins across WT and mutant FUS proteins, we performed MS-based proteomics using label-free quantitation (LFQ). A total of 4,954 unique proteins were identified and quantified across all 16 samples (4 replicates across 4 conditions). Significance Analysis of INTeractome (SAINT) analysis was performed determine whether putative interactions (prey) for each APEX2-FUS (bait) variant were valid (60). Prey with spurious interactions across all APEX2-FUS baits (*sensu* SAINT analysis; probability < 0.95) were eliminated from further analysis. Finally, the mean intensity of the control samples was subtracted from each sample intensity value for remaining prey proteins, leaving 3,349 proteins classified as putative interacting proteins in at least one sample.

Of the 3,349 proteins that met our filtering criteria, 3,229 (96.4%) were shared between all three groups suggesting substantial redundancy in binding partners between our three FUS variants (Supplementary Figure 2D). However, analysis from unsupervised hierarchical clustering analysis, heatmap analysis, and principal component analysis of each variant suggested that each variant enriched for the specific set of proteins (Figure 2C; Supplementary Figure 2B/CE). Given this, we reasoned that although the three variants enriched for many of the same proteins, the top proteins (i.e. the proteins most enriched for each variant) may be unique. As such, we specifically compared the top 10% most abundant proteins for each group (Figure 2A). We identified a total of 458 proteins in the top 10% of biotinylated proteins across the three variants. Unlike the full dataset of proteins (Supplementary 2D), only 197 proteins (43.0%) were shared between the three groups suggesting that each variant preferentially bound a different subset of proteins (Figure 2A). While APEX2-FUS PM had no unique proteins in the subset of enriched binding partner sharing 305 of its proteins (92.7%) with APEX2-FUS WT (Figure 2A), we identified 21 unique proteins in the top 10% most abundantly labeled proteins for APEX2-FUS WT and 105 unique proteins for APEX2-FUS P525L (Figure 2B). Taken together, these data suggest that while FUS PM interacts with a majority of the FUS WT binding partners, among the top 10% of proteins that interact with FUS PM, a subset exclusively interacts with pathogenic FUS P525L and not FUS WT. These interactions may impart novel functional characteristics to FUS PM that differ from FUS WT.

**Figure 2.**
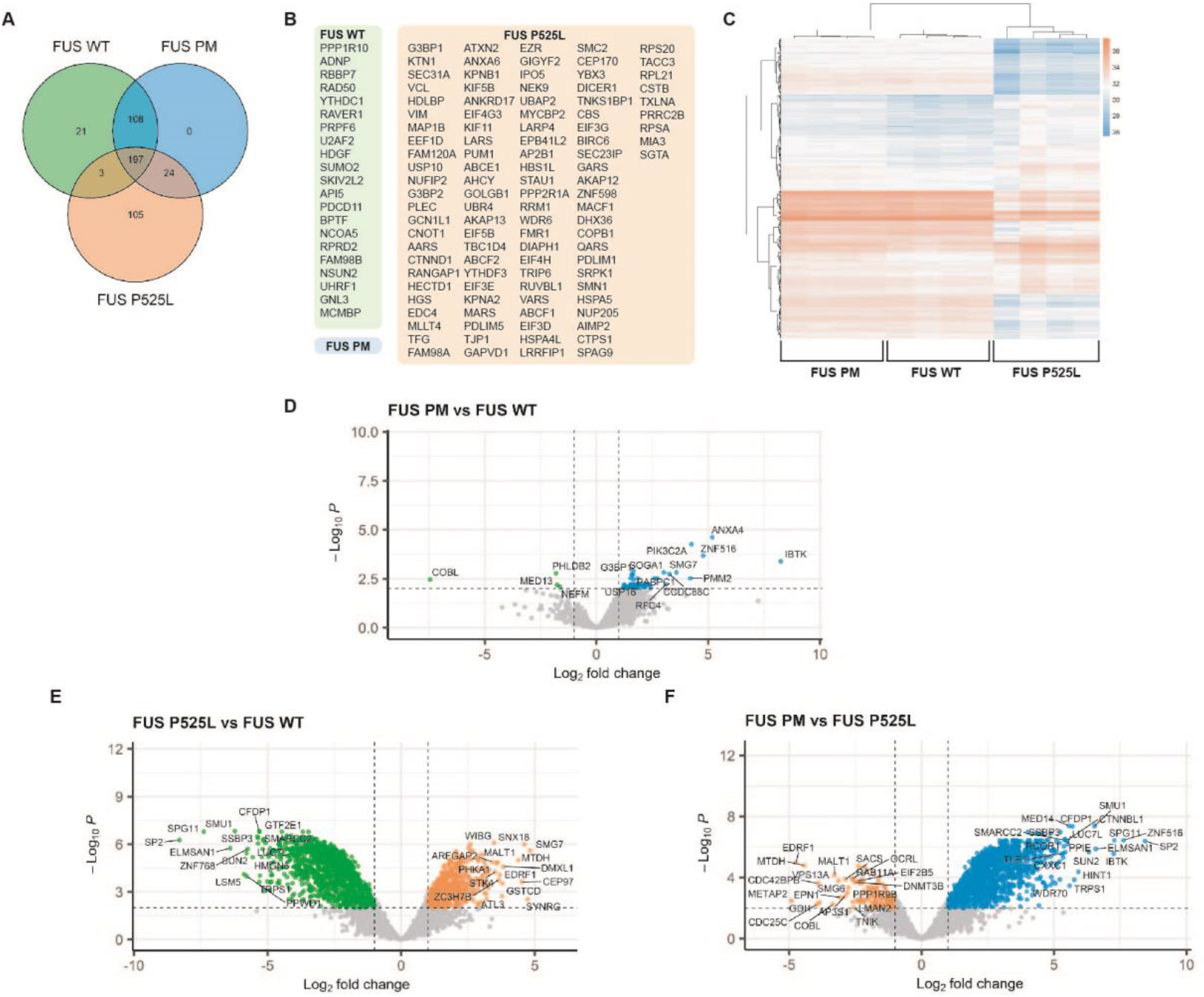
The FUS WT, FUS PM, and FUS P525L variants have unique interactome signatures. (A) Venn diagram of overlap of top 10% enriched proteins for the three FUS variant groups. (B) Proteins identified uniquely in each group are highlighted in colored boxes for three FUS variants. (C) Hierarchical clustering of samples based on the intensity profiles of the 10% of protein hits. Missing values are colored gray. (D) Volcano plot showing statistically significant enriched proteins identified between FUS WT and FUS PM. (E) Volcano plot showing statistically significant enriched proteins identified between FUS PM and FUS P525L. (F) Volcano plot showing statistically significant enriched proteins identified between FUS P525L and FUS WT.

Next, we compared the relative abundance of the interacting proteins between the FUS WT and FUS PM variants (Figure 2D), the FUS P525L and FUS WT variants (Figure 2E), and the FUS PM and FUS P525L variants (Figure 2F). For each comparison, we utilized a stringent cutoff of p<0.01 to produce a dataset of significantly enriched proteins for each variant. We then used Metascape to compare each differentially expressed gene set to the core ontologies (e.g. gene ontology (GO), KEGG processes, Reactome gene sets, canonical pathways and CORUM complexes) in order to gain insight into what biological processes or functional categories may be altered by each FUS variant (61) (Table 1). 53 proteins (1.6% of total identified proteins) differed between FUS WT and FUS PM, 181 proteins (5.4% of total identified proteins) differed between FUS PM and FUS P525L and 1590 proteins (47.5% of total identified proteins) differed between FUS WT and FUS P525L (Figure 2D/E/F). Of the 53 proteins differentially expressed in APEX2-FUS PM over APEX2-FUS WT, the top ontology categories are “mRNA catabolic process”, “translational assembly”, “stress granule assembly”, and “clathrin-mediated endocytosis” (Table 1). These functional categories occur in the cytoplasm suggesting FUS PM participates in more cytoplasmic pathways compared to FUS WT.

**Table 1.**
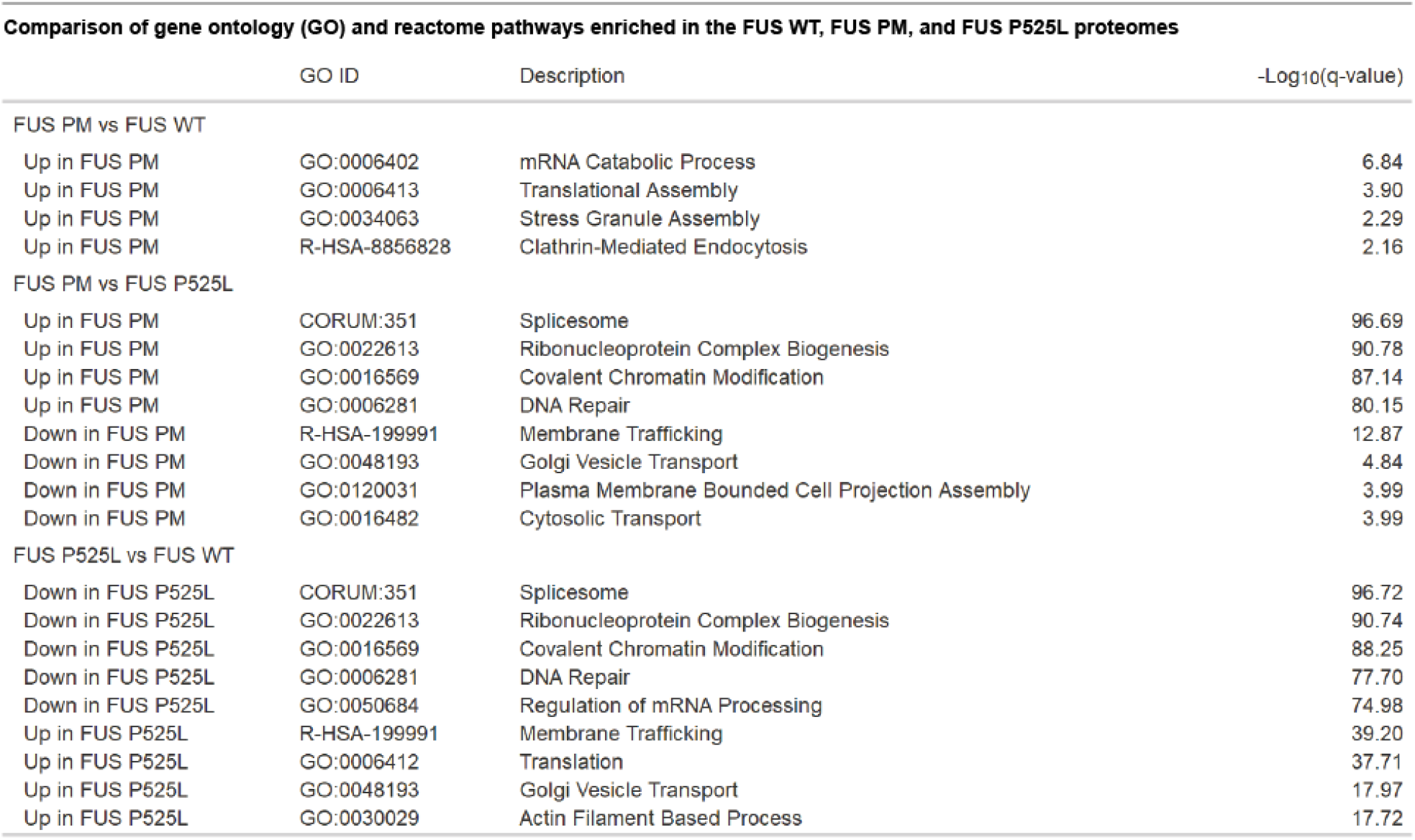
Comparison of gene ontology (GO) and reactome pathways enriched in the FUS WT, FUS PM, and FUS P525L interactomes. Table of statistically enriched gene ontology (GO) and reactome pathways generated using Metascape, a web-based platform designed to provide users a comprehensive annotation of provided gene list.

We also identified a subset of novel binding partners for FUS in our datasets. For the 4 proteins enriched in FUS WT over FUS PM we were unable to determine a categorical designation. These proteins were COBL, PHLDB2, MED13, and NEFM and are (Cordon-Bleu WH2 Repeat Protein, Pleckstrin homology-like domain family B member 2, Mediator of RNA polymerase II transcription subunit 13, and Neurofilament Medium Chain, respectively) all potentially novel binding partners for FUS WT. Furthermore, the top 4 enriched proteins for FUS PM compared to FUS WT were IBTK, PIK3C2A, ZNF516, ANXA4 (Inhibitor Of Bruton Tyrosine Kinase, Phosphatidylinositol-4-Phosphate 3-Kinase Catalytic Subunit Type 2 Alpha, Zinc Finger Protein 516, and Annexin A4, respectively) and are also potentially novel binding partners for FUS.

181 proteins were differentially enriched between APEX2-FUS PM and APEX2-FUS P525L (Figure 2E). Of these proteins, clustering analysis revealed FUS PM enriched for proteins associated with functions in the nucleus including “spliceosome”, “ribonucleoprotein complex biogenesis”, “covalent chromatin modification” and “DNA repair”. APEX2-FUS P525L variant enriched for pathways that occur in the cytoplasm including “membrane trafficking”, “Golgi vesicle transport”, “plasma membrane bounded cell projection” and “cytosolic transport” (Table 1). Lastly, we identified 1590 proteins differentially enriched between APEX2-FUS WT and APEX2-FUS P525L (Figure 2F). Of these proteins, clustering analysis revealed FUS WT enriched for proteins associated with the nuclear functions of “spliceosome”, “ribonucleoprotein complex biogenesis”, “covalent chromatin modification” and “DNA repair” while FUS P525L enriched for proteins associated with the cytoplasmic functions of “membrane trafficking”, “translation”, “Golgi vesicle transport” and “actin filament-based process” (Table 1).

Next, we constructed dot plots to clearly visualize the intensity and confidence of the protein interaction across each APEX2-FUS variant using the Prohits-viz software suite (62). We compared the binding partners identified in the top 4 significantly enriched ontology categories for FUS PM vs FUS WT (gene ontology or reactome) (Figure 3A/B/C/D). The binding intensity of the target proteins for FUS WT, FUS PM, and FUS P525L variants tended to occur as low, medium, and high, respectively. This observation compliments the original observation from the Venn diagram and the hierarchical cluster that FUS PM may exist in a middle state between FUS WT and FUS P525L function. A full list of dot plots for each identified ontology can be found in Supplementary Figure 3.

**Figure 3.**
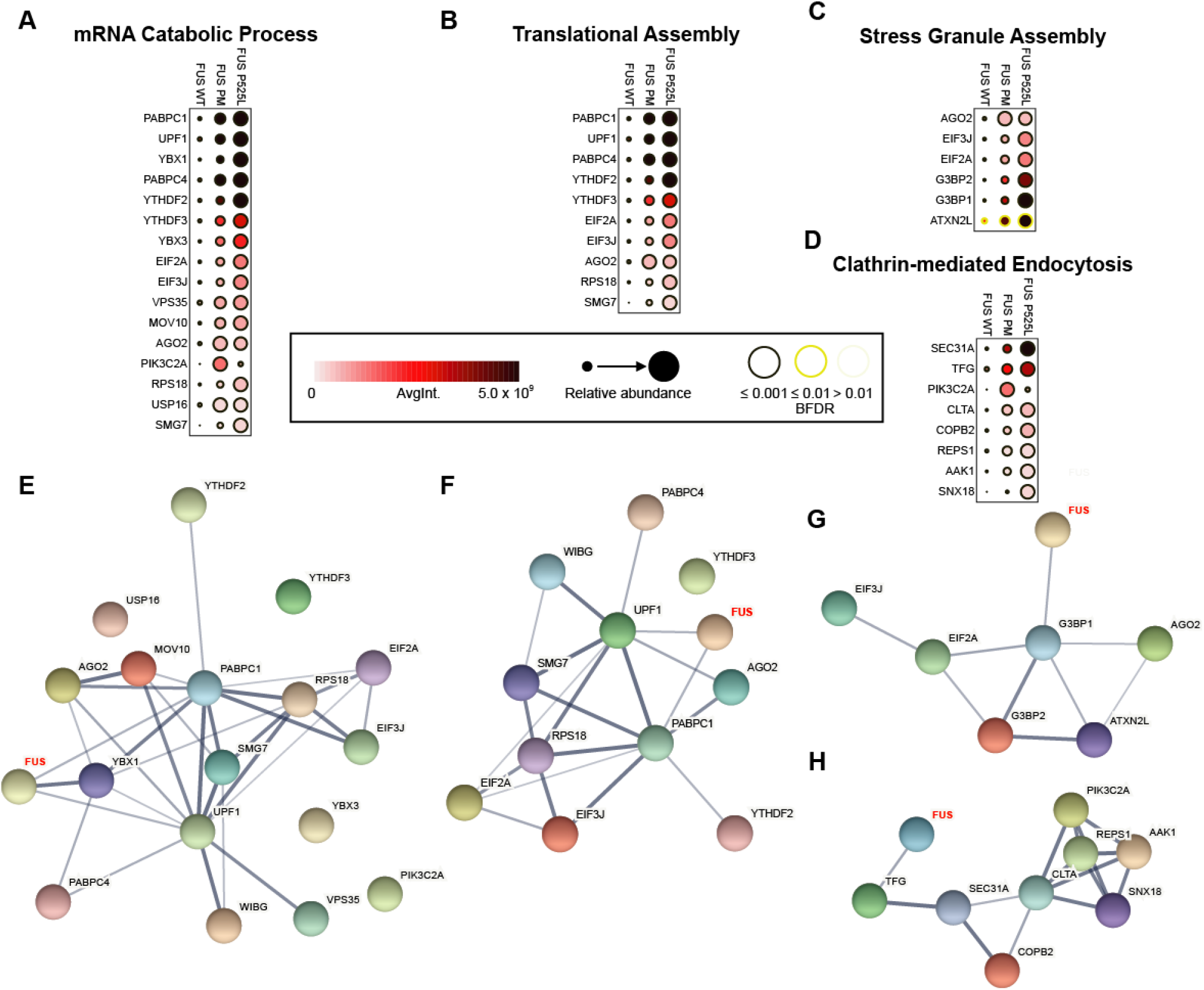
Visualization of protein hits for top gene ontology (GO) and reactome pathways reveals previously known and novel interaction partners of FUS variants. (A/B/C/D) Dot plot generated using ProHits-viz are a graphical representation of the relative binding intensity for the proteins mapped to (A) mRNA catabolic process, (B) translational assembly, (C) stress granule assembly, and (D) clathrin-mediated endocytosis GO term to the three FUS variants. (E/F/G/H) Protein interaction network for (E) mRNA catabolic process, (F) translational assembly, (G) stress granule assembly, and (H) clathrin-mediated endocytosis generated using String (version 11). Thickness of line between proteins indicates the strength of the empirical support for the interaction. FUS (in red) was added to network to demonstrate known binding partners.

Given that these GO terms were generated from gene sets of enriched proteins, we wanted to visualize the known interactions between FUS and the target genes of each gene set. We utilized the STRING database (version 11) to create an interaction network from each functional term (63) (Figure 3E/F/GH). The STRING algorithm is built from a curated list of known protein interactions to estimate how likely the interaction is true given the available evidence (termed confidence). The confidence for each interaction is shown by the thickness of the line between each protein. In these networks, we observed with high confidence that FUS interacts with some of the proteins in each network. Even so, there are few reports from previous studies indicating that FUS directly interacts with most of the proteins in each gene set. This may indicate that FUS WT interacts with more proteins in each interaction network than previously thought. Furthermore, this data suggests that N-terminal phosphorylation shifts the interaction landscape of FUS to interact with more proteins central to these functional categories. Leading us to ask, does FUS interact with the proteins identified in the gene sets? To answer this, we selected a subset of proteins (both previously identified as direct interactions and novel interactions) from the gene sets to validate using traditional biochemical approaches (immunoprecipitation and immunofluorescence): G3BP1, UPF1, MOV10, eIF2α, VPS35, PABPC1 (PABP1), and CLTA.

### Biochemical validation of FUS variant binding partners reveals novel interactions between FUS variants and APEX2 hits

We evaluated whether the FUS variants co-immunoprecipitated with the following selected endogenous targets: G3BP1, UPF1, MOV10, eIF2α, VPS35, PABPC1 (PABP1), and CLTA (Figure 4A). HEK293T cells were transfected with either Twin-Strep-tagged® FUS WT, FUS PM, or FUS P525L. All constructs were also GFP tagged at the N-terminus to allow visualization following transfection. We enriched for the strep-tagged FUS variants using Strep-Tactin®XT magnetic beads (IP) and western blotted for the potential endogenous binding partners (IB) (Figure 4A). As members of the FET family of proteins, EWS and TAF-15 are known binding partners of FUS and were used as a positive control for interaction (Figure 4A, green bar) (dot plot for EWS and TAF15 in Supplementary Figure 3) (64). We were able to replicate that FUS WT and FUS P525L co-IP with UPF1, PABP1, G3BP1, and eIF2α as previously reported (Figure 4A, blue bar) (53,65–67).

**Figure 4.**
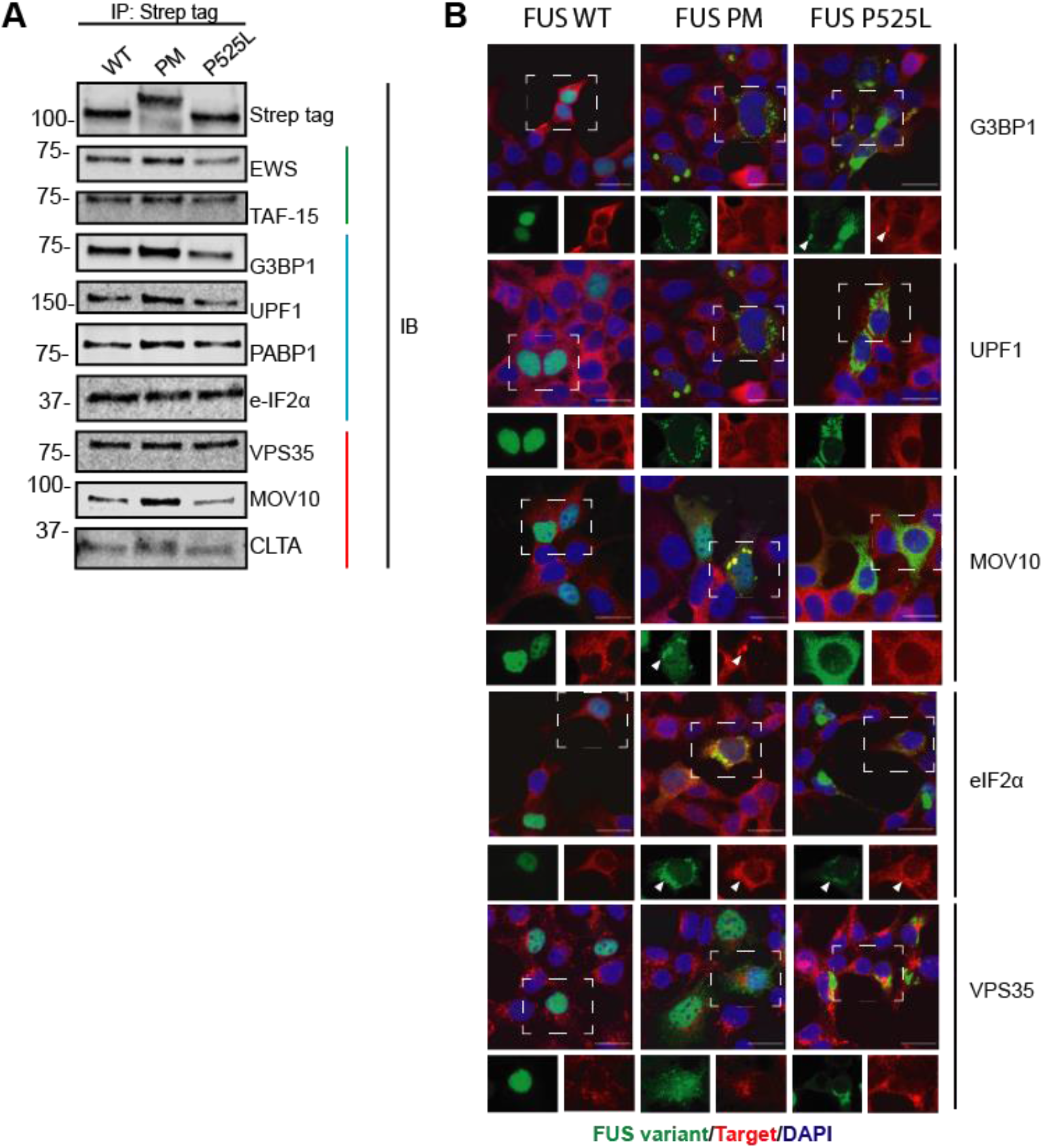
Verification of the interaction between select targets and FUS variants. (A) Immunoprecipitations (IP) for strep tag were performed on HEK293T cells expressing GFP-tagged FUS WT, FUS PM, and FUS P525L. Enriched lysate was immunoblotted (IB) for listed targets. (B) Immunofluorescence (IF) images show general localization patterns for a select number of targets. Co-localization of targets with FUS punctate is highlighted by white carrot. FUS variants are in green, targets are in red, DAPI is in blue. Scale bar represents 20µm.

Next, we asked if FUS PM FUS PM interacted with proteins in a similar manner to N-terminally phosphorylated FUS, which occurs in cells following DNA damage. To do this, we performed a co-IP of FUS from HEK293T cells treated with calicheamicin γ1 (CLM), a known inducer of N-terminal FUS phosphorylation (47,49,50). We found endogenous UPF1 preferentially co-IP’d with CLM-induced phosphorylated FUS (Supplementary Figure 4). Lastly, we confirmed the interaction of three novel binding partners, VPS35, MOV10 and CLTA, to our three FUS variants (Figure 4A, red bar). This is the first report that FUS PM interacts with any of these proteins.

Given that the three FUS variants are enriched in different cellular compartments (Figure 1H), we performed immunofluorescent staining for a subset of the top proteins to determine the spatial localization of the binding partners with the FUS variants (Figure 4B; PABP1, EWS and TAF15 not shown). We expressed Twin-Strep-tagged® FUS variants in HEK293T and then co-stained for the endogenous target proteins. As expected, FUS WT was enriched in the nuclear compartment while FUS PM and FUS P525L localized to the cytoplasm. The endogenous target proteins localized to cytoplasm. Given this, we saw spatial overlap of the endogenous target proteins with FUS PM and FUS P525L. For G3BP1 and MOV10, this overlap, at times, occurred in large puncta (Figure 4B, white arrow). Thus, our APEX2 generated dataset shows robust agreement with our biochemical validation.

### The steady-state level of ATF3 transcripts is increased while global protein translation is enhanced in the presence of FUS PM

Following validation of protein targets identified by APEX2, we set out to test whether the functional pathways suggested by our enrichment analysis were affected by the expression of a given FUS variant. We utilized 4 N-terminally GFP/Twin-Strep-tagged® FUS constructs: 1) wild-type human FUS (WT), 2) human FUS where the 12 serine/threonine’s phosphorylated by DNA-PK are substituted with alanine (Ala sub), 3) human FUS where the 12 serine/threonine’s phosphorylated by DNA-PK has been substituted with the negatively charged aspartate (PM), 4) human FUS truncated at exon 15 (delta 15). We utilized the delta 15 truncation mutant as a proxy for the P525L mutation and the pcDNA3.1 empty vector (EV) as a control.

We specifically focused on the pathways enhanced by FUS PM expression. The highest enriched ontology category for FUS PM over FUS WT was “mRNA catabolic process”, defined as the reactions and pathways associated with the breakdown of mRNA (Table 1). As an RNA/DNA binding protein, FUS expression has been shown to regulate ∼700 mRNA transcripts related to the regulation of transcription, RNA processing, and cellular stress response (68). Expression of ALS-linked mutations in FUS can shift the global transcriptome (12, 69). Specifically, a previous study reported that degradation of certain mRNA transcripts is increased following expression of the ALS-linked mutant FUS P525L (53). We also observed a positive interaction between the FUS variants and UPF1 and PABP1, both major mediators of the mRNA decay pathway, nonsense mediated decay (NMD). As such, we asked whether expression of FUS PM altered the steady state levels of specific mRNA transcripts regulated by NMD (70). We designed a quantitative PCR (qPCR) protocol to measure the total levels of the stress-related mRNA targets ATF3, ATF4, and TBL2 (Supplementary Table 2). While total mRNA level for UPF1 and FUS were not significantly different between FUS variants, we saw a significant increase in ATF3 mRNA levels in HEK293T cells expressing PM compared to EV, WT, and delta 15 (p=0.0034, p=0.0357, and p=0.0008, respectively) (Figure 5B). Further, we observed a trend for an increase in ATF4 mRNA in HEK293T cells expressing delta 15 but not PM (EV vs. delta 15, p=0.3721; WT vs. delta 15, p=0.3994). We saw no difference in TBL2 mRNA levels following expression of FUS variants. Taken together, this data suggests that the steady state levels of certain transcripts are increased by FUS PM expression.

**Figure 5.**
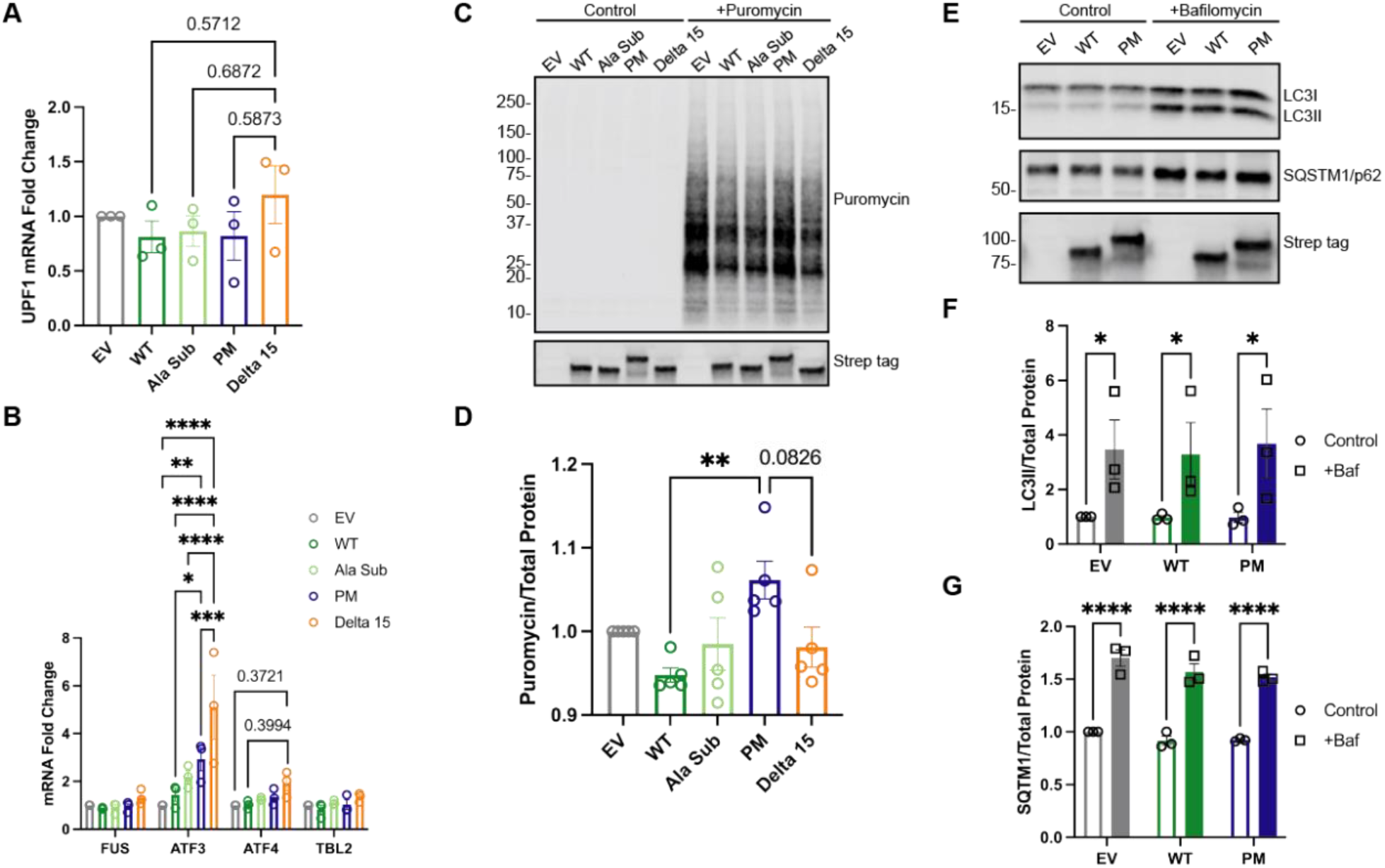
Biochemical validation of GO pathways show alternations in nonsense mediated decay and translation independent of clathrin-mediated endocytosis between the FUS variants. (A) Level of UPF1 mRNA was quantified by qPCR using the ΔΔ cycle threshold (ΔΔCT) method and then the fold change was calculated against the empty vector control (EV). (B) Level of various stress related targets of mRNA decay were quantified by qPCR using the ΔΔ cycle threshold (ΔΔCT) method and then the fold change was calculated against the empty vector control (EV). (C) Representative immunoblot of SUnSET assay measuring the incorporation of puromycin into growing polypeptide chains during translation. Control cells received HEPES buffer without puromycin for 30 minutes. (D) Quantification of immunoblot in (C). Error bars indicates mean ± SEM (n=5). Statistical significance was calculated by one-way ANOVA. (E) Representative immunoblot for markers of autophagosome flux, LC3I/II and SQSTM1/p62. (F) Quantification of immunoblot in (C). Error bars indicates mean ± SEM (n=3). Statistical significance was calculated by two-way ANOVA.

Next, we set out to examine whether expression of FUS PM affected mRNA translation, the closely linked functional process. NMD is thought to be tightly coupled to translation because 1) translation requires multiple NMD factors, 2) phosphorylated UPF1 suppresses translational initiation, and 3) re-initiation of translation downstream of the premature termination codon can prevent NMD (71). Expression of FUS P525L was previously reported to decrease global protein translation (53). Taken together, with the fact that the next highest enriched functional pathway was “translational assembly”, we utilized the SUrface SEnsing of Translation (SUnSET) assay to compare the amount of global protein synthesis between the FUS variants (72) (Figure 5C). We saw a significant increase in the amount of protein synthesis in HEK293T cells expressing FUS PM compared to FUS WT (p=0.0074) (Figure 5D). Furthermore, we saw a trend toward a decrease in protein synthesis between PM and delta 15 (p=0.0826) (Figure 5D). As such, protein translation is unchanged by delta 15 expression and enhanced by FUS PM expression compared to FUS WT.

Lastly, we examined the amount of autophagosome formation as a proxy for “clathrin-mediated endocytosis”. Clathrin-coated vesicles form the precursor phagophores and blocking clathrin-dependent endocytosis leads to a decrease in autophagosome formation (73). Further, two of our protein hits, CLTA and VPS35, are important for autophagosome formation (74). Autophagosomes are double membrane vesicles that are integral to macroautophagy as they sequester cellular components and eventually fuse with acidic lysosomes to form autolysomes and degrade engulfed material (75, 76). We utilized an autophagic assay where we treated cells with bafilomycin (Baf), an inhibitor of the lysosomal V-ATPase in order to block the fusion of autophagosomes leading to a build-up autophagosomes (77). There was no difference in the levels of the autophagosome markers, LC3II and SQSTM1/p62, following expression of FUS variants, before or after Baf treatment. Overall, these data suggest that while FUS PM expression affects early mRNA translation and regulation, it does not affect the total amount of autophagosomes or autophagosome flux.

## Discussion

Proteomic analysis is a powerful tool that has revealed how pathogenic ALS-linked mutations (e.g. FUS P525L and R495X) may lead to pathology (14,52,59). Past studies have utilized both targeted immunoprecipitations and whole cell analysis to map the proteome of cells expressing various toxic ALS-linked FUS mutations (i.e. P525L, R495X) (52,53,59). Interactome level changes have been mapped for wild-type (WT) and ALS-linked R521G FUS (14). Furthermore, a recent study looked specifically at the interactome level changes in pathologically relevant droplets purified from WT and P525L FUS expressing cell lysates (54). While these past studies provide some insight into the role that *FUS* mutation can have on protein-protein interactions, pathogenic FUS mutations only account for 4% of ALS cases and a handful of FTD cases (33,34,78,79). Thus, these previous studies do not address how non-genetic triggers of FUS pathology, such as post-translational modifications, may shift FUS function. Previous non-genetic models demonstrate that cytoplasmic accumulation of FUS can be triggered by other, non-genetic mechanisms including loss of transportin-1/FUS interaction, cellular stressors, and/or altered post-translational modifications (15,35-40,49). Although methylation state of FUS can be shifted in ALS/FTD-FUS post-mortem tissue, causative genetic mutations, or cellular stressors, have not been identified to explain this phenomenon (37, 80). Unlike methylation, our lab has shown that a biologically relevant stressor, double strand DNA breaks (DSBs), triggers the DNA dependent protein kinase (DNA-PK) to phosphorylate FUS at 12 key S/T_Q residues in the N-terminal SYGQ-low complexity domain (45-47,49,50). Phosphorylated FUS (p-FUS) then accumulates in the cytoplasm of the cell (49, 81). While previous studies have examined how DNA-PK mediated N-terminal phosphorylation of FUS may shift the structure of the N-terminus of FUS towards a more disordered state *in vitro*, none have determined whether phosphorylation at these residues alters the function of FUS in cells (45, 46). In this study, we investigated whether N-terminal phosphorylation at these 12 key residues shifts the FUS protein interactome and its cellular functions. We utilized the APEX2 system in combination with label-free proteomic analysis to investigate the role of N-terminal phosphorylation in the SYGQ-rich low complexity domain on FUS function. Overall, this study is the first to map changes in the FUS protein interactome associated with a PTM.

The first question we aimed to address was whether the labeled protein hits of FUS PM overlap more with homeostatic FUS WT or pathogenic FUS P525L. From the 3,349 proteins we identified in our study, 96.4% were shared between all three FUS variants (Figure 2A/C). This suggests that the pathogenic FUS P525L and the DSB-associated FUS PM variants can interact with the majority of FUS WT targets (Figure 2A). This is surprising as pathogenic variants of another ALS/FTD linked protein, TDP-43, have been shown to interact with a large proportion of novel binding partners compared to wild-type TDP-43 (82, 83). As such functional changes seen in pathogenic FUS P525L and the DSB-associated FUS PM variants may not be due to the development of novel protein interactions but instead are related to changes in the strength of interaction partners. For instance, methylation of key C-terminal residues in the RGG3 domain greatly shifts the strength of the interaction between FUS and its major nuclear import protein, transportin-1 (TNPO1) (37, 84). In line with this, our data supports the idea that FUS pathology is not due to a general loss of FUS interaction with target proteins since pathogenic FUS P525L interacted with most FUS WT target proteins (25, 85). These findings suggest that pathogenesis may be due to changes in the strength of FUS interactions with other proteins.

To examine whether the strength of interactions between the FUS variants and protein hits differed, we focused on the top 10% most enriched protein hits for each variant and looked at the overlap of each group (Figure 2A). Each sample clearly separated into three distinct groups, with FUS WT and FUS PM overlapping more than FUS P525L (Figure 2A/C). This distribution suggests that while a majority of the protein interaction network is shared between the three groups, the datasets from FUS WT and FUS PM share more in common with each other than FUS P525L. If the protein binding partners of FUS PM mirror FUS WT more than FUS P525L, does this indicate that expression of FUS PM is not disruptive the FUS interactome? To answer this question, we utilized differential expression analysis to directly examine the relative differences in abundance between our three groups. We saw that the comparison of FUS WT and FUS P525L exhibited the highest number of differentially expressed proteins followed by the comparison of FUS PM and FUS P525L (Figures 2D/E/F). Likewise, FUS PM also enriched for a subset of proteins over FUS WT, suggesting that FUS PM may participate in biology processes in a manner divergent from that of FUS WT.

We took advantage of the list of differentially enriched genes between our groups to understand whether FUS function was affected by FUS PM expression. Past studies have demonstrated that expression of FUS P525L leads to functional changes in ontological pathways including altered translation, altered splicing, and dysregulated chromatin (52,53,86–88). In line with these past studies, our APEX2-FUS P525L dataset was enriched for both cytoplasmic functional terms (“translation”) and structural terms (“actin filament-based process”), while depleted for nuclear terms related to mRNA (“spliceosome” and “regulation of mRNA processing”) and DNA processes (“covalent chromatin modification” and “DNA repair”). As such, APEX2-FUS P525L proximity biotinylated proteins tended to be localized to the cytoplasm, suggesting cytoplasmic functional pathways may be altered by FUS P525L expression (Figure 1H) (28). Our FUS WT vs FUS P525L dataset agrees with previous functional studies demonstrating that nucleocytoplasmic shuttling is an important mediator of FUS function.

The identified subgroup of enriched ontology terms for FUS PM over FUS WT were “mRNA catabolic process”, “translational assembly”, “stress granule assembly” and “clathrin-mediated endocytosis”. These terms covered primarily cytoplasmic functions consistent with the observation that FUS PM accumulates in the cytoplasm over FUS WT (Figure 1D). Even so, the role of N-terminal phosphorylation in pathology is a debated topic. Other studies report N-terminal phosphorylation reduces the propensity of FUS to aggregate *in vitro*, thereby supporting a model where phosphorylation may be protective against cytoplasmic FUS-mediated toxicity (45, 50). In contrast, we provide evidence that N-terminal phosphorylation instead promotes the formation of FUS aggregates (Supplementary Figure 5). Aggregation of FUS, independent of a pathogenic genetic mutation, may itself be sufficient to induce neurodegeneration (32). As such, this may suggest that aggregates of N-terminally phosphorylated FUS may induce cellular toxicity. Future studies will need to investigate the role these aggregates have in cellular heath.

Next, we utilized Prohits-viz to directly compare the abundance of the binding hits identified for these four ontology terms between each FUS variants (Figure 3). From this, we were able to visualize a multitude of proteins that overlap between ontology categories. We used this data along with the STRING interaction database to identify a subset of proteins from each ontology term that were either 1) previously identified binding partners for FUS WT (G3BP1, UPF1, PABP1, eIF2α) or 2) novel binding partners (VPS35, MOV10, CLTA) (14,41,53,65,67). As anticipated by the APEX2 datasets, we were able to confirm the interaction between all three FUS variants and the above targets utilizing two different methods: immunoprecipitation and immunofluorescence (Figure 4A/B).

First, we confirmed that the GFP-tagged FUS PM and FUS P525L localized to the cytoplasm (Figure 4B). Further, FUS PM and FUS P525L co-localized in the cytoplasm with the target proteins (G3BP1, UPF1, MOV10, eIF2α and VPS35 (Figure 4B). Even though FUS WT did not form distinct puncta or aggregates with these target proteins, it should be noted that FUS is a nucleocytoplasmic protein that shuttles between these two cellular compartments (20, 89). Therefore, while FUS WT accumulation into in the nuclear compartment is easily visualized through immunofluorescent staining, a significant portion of the protein is cytoplasmic (Figure 1C). As such, we confirmed that all three FUS variant co-IP’d with previously identified binding partners (EWS, TAF-15, G3BP1, UPF1, PABP1, and eIF2α). We also confirmed novel interactions between all three FUS variants and VPS35, MOV10, and CLTA. VPS35, MOV10, and CLTA may have been previously linked to ALS/FTLD pathology. VPS35 is a key component of the retromer trafficking complex and is highly expressed in pyramidal neurons, a key cellular target in FTLD-mediated pathology (74). MOV10 is a member of the SF-1 RNA helicase family related to UPF1 and a component of the RNA-induced silencing complex (RISC) (90). Exogenous expression of MOV10 was shown to ameliorate cell death in a TDP-43 model of ALS pathology (91). Decreased expression of the endocytic protein, CLTA, has been reported as a potential general marker of early endocytic dysfunction in neurodegeneration (92). While all three of these proteins have been previously linked to FTD/ALS pathology, no study has directly linked FUS binding to these targets. The positive validation of these targets may open new avenues to explore FUS dysfunction.

Next, we set out to determine the extent that FUS PM expression affected functional pathways suggested by the proximity-labeling. Alterations in mRNA catabolic processing have been strongly linked to both ALS and FTD. One such process is nonsense mediated decay (NMD). NMD is a major cellular mechanism responsible for mRNA quality control by surveilling mRNA for premature termination codons (71, 93). UPF1 and PABP1, two proteins differentially enriched in FUS PM over FUS WT, act as opposing forces mediating the degradation/stabilization of NMD-sensitive mRNAs (94). A recent report found that NMD was inhibited in a C9orf72-model of FTD pathology, indicating that NMD dysfunction could be a common finding across the ALS/FTD spectrum (95). Overexpression of UPF1 in a model of FTD ameliorated toxicity in a model of ALS, suggesting enhancing NMD may be beneficial (96). In contrast, another report found that an ALS-linked FUS mutant enhanced NMD decay of targeted transcripts (53). What might explain these discrepancies? One possibility is that past studies utilized model systems derived from different species. Studies that found diminished NMD were performed in human-derived models or using an *in vivo* mouse model of FUS pathology, while the study that shows enhanced NMD was done in an immortalized mouse cell line (53,95,97). Recently, we reported that mouse cells do not recapitulate DSB-mediated N-terminal phosphorylation of FUS (49), raising the possibility that FUS-mediated regulation of NMD is also not accurately recapitulated in mouse cells. To avoid these species-specific differences, we measured the steady-state levels of known targets of NMD using a qPCR assay in human HEK293T cells. We found that mRNA transcript levels of ATF3, but not ATF4, are significantly increased following expression of FUS PM and truncated FUS delta 15 (Figure 5B). These data suggest that expression of FUS PM may shift the steady-state levels of certain mRNA transcripts. Future studies will need to explore if NMD processes are responsible for this shift.

What might be causing this divergence in steady state ATF3 but not ATF4 transcript levels? Various cellular stressors such as the production of reactive oxygen species or ER stress leads to upregulation of ATF3 and ATF4 (98). ATF3 is a stress-induced transcriptional activator associated with binding genomic sites related to cellular stress (99). In parallel, expression of ATF4 leads to ATP depletion, oxidative stress, and cell death (83). Interestingly, upregulation of ATF4 occurs first, before directly inducing the expression of ATF3 and other downstream transcriptional regulators. Given that we only assayed one time point, it is possible that while the 48-hour time point captures the change in total transcripts levels for ATF3, it may be too late to detect appreciable changes in ATF4 transcript levels (98, 100). Furthermore, we did not detect altered transcript levels for two other targets of NMD (Figure 5A/B) (98, 101). Thus, expression of FUS PM expression may target specific mRNA transcripts. Consistent with this idea, previous studies have shown that not all perturbations to the mRNA decay pathways equally affect transcript expression. For instance, depletion of NMD factor UPF2 enhanced ATF3 but not TBL2 mRNA transcript levels (101). Alternatively, induction of ATF3 mRNA following expression of FUS PM and delta 15 may reflect the role of ATF3 as a stress-responsive transcription factor (102). Future studies should investigate the role FUS phosphorylation on stress response pathways and the specificity of mRNA catabolic suppression on other transcripts.

FUS function is closely linked to regulation of mRNA translation (52,53,103–105). In line with this, we saw that expression of FUS PM enhanced protein translation compared to FUS WT (Figure 5D). Interestingly, while we saw a trend, we did not find a significant change in protein synthesis following FUS delta 15 expression (Figure 5D). Cytoplasmically localized ribonucleoprotein complexes (RNP) granules containing FUS, wild-type or an ALS mutant, have been reported to participate in active protein translation (105). Accordingly, FUS PM and FUS P525L in the cytoplasm may enhance protein translation through a similar mechanism. It should be noted that while the SUnSET is thought to reliably measure protein translation it does have some limitations: 1) it measures relative rates of synthesis and is unable to capture the absolute changes and 2) differences in the amount of free puromycin between samples may alter puromycin uptake (72). Therefore, future studies should compare multiple methods of quantifying protein synthesis.

Lastly, we examined how expression of FUS PM may impact autophagy through autophagosome formation. Lysosome-mediated autophagy is a multi-stage process involving multiple cellular components. In this process, autophagosomes are an integral part of the autophagy cascade where they begin as phagophores that expand into autophagosomes and fuse with endosomes and lysosomes to allow degradation of the compartment contents (73). Dysfunctional autophagosome formation and other aspects of the autophagy-lysosome pathway has been widely reported in ALS and FTD (106). In this study, we idented CLTA as a binding partner for FUS, which is involved autophagosome formation. However, we did not detect any difference in the levels of two markers of autophagosomes following FUS WT and FUS PM expression, suggesting autophagosome formation is not affected (Figures 5F/G) (107). Nonetheless, it remains possible that phosphorylation of FUS, or expression of pathogenic *FUS* mutations, affects autophagy and related pathways (e.g. autophagic flux, lysosome health, fusion, endocytosis) (106, 108). Future studies should examine whether other parts of the clathrin-mediated endocytic pathway are affected by expression of FUS PM.

In conclusion, we report the first study examining whether a post-translational modification, N-terminal phosphorylation, affects the FUS proteome. Using the APEX2 system, we identified a robust dataset of novel protein partners for FUS WT, FUS P525L, and a mimetic of N-terminal phosphorylation of FUS. We provide evidence that expression of phosphorylated FUS may impact cellular function by enhancing translation and suppressing mRNA degradation. These findings also shed light on fruitful avenues for future investigation. Future studies should examine how post-translational modifications of FUS regulate protein function within the cell and how non-genetic factors influence processes underlying disease. The discovery that phosphorylated FUS may play a unique role in the mRNA homeostasis provides valuable insights into what functions may be dysregulated in the pathological cascades of ALS and FTD.

## Experimental Procedures

### Plasmid creation

APEX2-FUS plasmids, maps, and sequences generated in this study are deposited in Addgene. The DNA sequences for the APEX2-FUS variants were designed *in silico* then codon optimized and custom synthesized by GenScript. The amino acid sequence for the engineered APEX2 was taken from Addgene plasmid #212574. The wild-type FUS sequence was taken from NCBI reference sequence RNA-binding protein FUS isoform 1 [Homo sapiens] (NP_004951.1). A Twin-Strep-tag® was added to the N-terminus of the APEX2 sequence. A linker region (GGGS)^3^ was inserted at the end of APEX2 followed by the FUS sequence. Synthetic APEX2-FUS gene constructs were designed to add a 5’ BamHI restriction digestion site (GGATCC) followed by a Kozak sequence (GCCACC) before the ATG start codon of APEX2, a 3’ stop codon (TAG) and an ending with a XhoI restriction digestion site (CTCGAG). Following synthesis, the APEX2-FUS WT fusion protein was inserted into the pcDNA3.1/Hygro(+) vector using a BamHI/XhoI cloning strategy. The APEX2-FUS P525L and APEX2-FUS PM constructs was generated from the donor APEX2-FUS WT construct by express mutagenesis through GenScript.

The GFP tagged FUS variants were designed by adding EGFP to the N-terminus of the previously described FUS variants in Deng et al. (47). In brief, the FUS variants (WT, Ala sub, PM, and delta 15) were synthesized and ligated into pcDNA3.1(+) Hygro by GeneArt (ThermoFisher Scientific). These constructs were then digested at NheI/HindIII sites upstream of the FUS sequence. EGFP was PCR amplified to introduce an NheI restriction site at the 5’ end and a HindIII site at the 3’ end. The EGFP was then digested and ligated into each construct. The primers used to generate EGFP were: GFP.Nhe.Sense (CACTATAGGGAGACCCAAGCTGGCTAGCgccaccATGGTGAGCAAGGGCGAGGAG CTG) and GFP.Hind.Antisense: (GGGACCAGGCGCTCATGGTGGCAAGCTTCTTGTACAGCTCGTCCATGCCGAG).

The GFP tagged FUS P525L variant was created by site directed mutagenesis on the GFP tagged FUS WT construct using the QuikChange II XL Site-Directed Mutagenesis Kit (Agilent; 200521). The primers used to generate the construct were: P525L_Sense (gacagaagagagaggctctactgactcgagtct) P525L_Antisense (agactcgagtcagtagagcctctctcttctgtc)

All constructs were verified using DNA sequencing, restriction digests, and/or PCR amplification. The full DNA sequence for each synthesized sequence can be found in Supplemental Table 1

### Cell culture

Human embryonic kidney cells (HEK293T, ATCC) were cultured in DMEM medium supplemented with 10% fetal bovine serum (FBS, Atlanta Biological) and 1% Pen/Strep (Gibco). Cells were maintained at 37°C with 5% CO_2_.

### Cell transfection and APEX2-mediated biotinylation

HEK293T cells were seeded onto a poly-L-lysine coated 10-cm cell culture grade dish and cultured for 2 days prior to transfection. Cells were transfected at ∼60% confluency with 2.5 µg of the appropriate DNA construct using the TransIT-LT1 Transfection Reagent (Mirus; MIR2300) and cultured for an additional 2 days. At ∼48 hours post transfection, 500 µM biotinyl tyramide (biotin phenol) (Tocris; #6241) supplemented in DMEM media with 10% FBS/1% Pen/Strep was added to all experimental plates except for the non-transfected control plates. Labeling was initiated after 30 minutes by adding H_2_O_2_ (1 mM final concentration) for 1 minute. The labeling reaction was quenched by aspirating media from the plate and immediately rinsing three times with the quenching solution: 5 mM trolox ((+/-)-6-Hydroxy-2,5,7,8-tetramethylchromane-2-carboxylic acid, Sigma; 238813), 10 mM sodium L-ascorbate (Sigma; A4034) and 10 mM sodium azide in PBS supplemented with 1x phenylmethylsulfonyl fluoride (PMSF), a serine protease inhibitor. Cells were then incubated on ice in fresh quenching solution four times for 5 minutes each. Following the last wash, the quenching solution was aspirated off and 600 µl cold lysis buffer (50 mM Tris, 150 mM NaCl, 0.4% SDS, 0.5% sodium deoxycholate, 1% Triton X-100, 10 mM sodium azide, 10 mM sodium ascorbate, and 5 mM Trolox) supplemented with 1x Halt protease/phosphatase inhibitor (ThermoFisher; 78446) was added to each plate. Samples were collected with cell scrapers into Protein lo-bind tubes (Eppendorf) and sonicated 2x on ice (25 amplitude: 10 seconds total on ice, 2 seconds on/2 seconds off). Samples were cleared by centrifugation at 16,500xg for 10 minutes at 4°C and supernatant was collected into fresh protein lo-bind tubes. 540 µl of pre-chilled 50 mM Tris pH=7.4 was added to wash each pellet and samples were spun at 16,500xg for 10 minutes at 4°C. Supernatant was collected and combined to previous samples and samples were stored at −80°C. Protein concentration was assayed using RC DC protein assay (Bio-Rad; 5000121).

### Streptavidin-based purification of biotinylated targets

For affinity purification, 240 µl of NanoLINK Streptavidin Magnetic Beads (TriLink Biotechnologies; M-1002) were washed 3x in 1x tris buffered saline (TBS) containing 0.1% tween-20. 1.8 mg of total protein was then added onto washed beads and allowed to incubate overnight at 4°C with mixing. Beads were then collected against a magnetic stand and the supernatant was set aside for future analysis (termed flow-through). Beads were then washed in wash buffer 1 (50 mM Tris, 150 mM NaCl, 0.4% SDS, 0.5% sodium deoxycholate, and 1% Triton X-100) and gently mixed with rotation for 5 minutes at room temperature. Supernatant was discarded. Beads were then washed in wash buffer 2 (2% SDS in 50 mM Tris HCl, pH 7.4) and gently mixed with rotation for 5 minutes at room temperature. Supernatant was discarded. Beads were then washed 2x in wash buffer 1 with rotation for 5 minutes at room temperature. 10% of bead slurry from each sample was set aside for future analysis (termed elution). Remaining beads were then washed 4x in 1x phosphate buffered saline (PBS) and stored at −20°C.

### On-bead digestion and label-free mass spectrometry

8 M urea was added to the beads and the mixture was then treated with 1 mM dithiothreitol (DTT) at room temperature for 30 minutes, followed by 5 mM iodoacetimide (IAA) at room temperature for 30 minutes in the dark. Proteins were digested with 0.5 µg of lysyl endopeptidase (Wako) at room temperature for 4 hours and were further digested overnight with 1 µg trypsin (Promega) at room temperature. Resulting peptides were desalted with HLB column (Waters) and were dried under vacuum.

### Mass Spectrometry

The data acquisition by LC-MS/MS was adapted from a published procedure (109). Derived peptides were resuspended in the loading buffer (0.1% trifluoroacetic acid, TFA). Peptide mixtures were separated on a self-packed C18 (1.9 µm, Dr. Maisch, Germany) fused silica column (50 cm x 75 µm internal diameter (ID); New Objective) attached to an EASY-nLC™ 1200 system and were monitored on a Q-Exactive Plus Hybrid Quadrupole-Orbitrap Mass Spectrometer (ThermoFisher Scientific). Elution was performed over a 106 min gradient at a rate of 300 nL/min (buffer A: 0.1% formic acid in water, buffer B: 0.1 % formic acid in acetonitrile): The gradient started with 1% buffer B and went to 7% in 1 minute, then increased from 7% to 40% in 105 minutes, then to 99% within 5 minutes and finally staying at 99% for 9 minutes. The mass spectrometer cycle was programmed to collect one full MS scan followed by 20 data dependent MS/MS scans. The MS scans (350-1500 m/z range, 3 x 10^6^ AGC target, 100 ms maximum ion time) were collected at a resolution of 70,000 at m/z 200 in profile mode. The HCD MS/MS spectra (2 m/z isolation width, 28% collision energy, 1 x 10^5^ AGC target, 50 ms maximum ion time) were acquired at a resolution of 17,500 at m/z 200. Dynamic exclusion was set to exclude previously sequenced precursor ions for 30 seconds within a 10 ppm window. Precursor ions with +1, and +8 or higher charge states were excluded from sequencing.

### Proteomic Data Processing

#### Raw Data Processing

Raw files were processed by MaxQuant with default parameters for label-free quantification (110). MaxQuant employs the proprietary MaxLFQ algorithm for LFQ. Quantification was performed using razor and unique peptides, including those modified by acetylation (protein N-terminal), oxidation (Met) and deamidation (NQ). Spectra were searched against the Human Uniprot database (90,300 target sequences). The resulting data with intensity scores were run through the Significance Analysis of INTeractome (SAINT) software (version 2.5) to identify and remove proteins that were unlikely to be true bait-prey interactions (60). This was performed by comparing protein intensity values in the negative control condition to the corresponding intensity values in the samples. Proteins with less than 95% probability to be significantly different from the negative control in all samples were removed. The mean intensity values of control were subtracted from each sample intensity value for the remaining proteins.

#### Statistical Analysis

The resulting protein groups information was read in R and analyzed using Proteus to determine differentially expressed proteins between groups (111). Label-free quantitation (LFQ) intensities of each sample were log_2_ transformed and compared using a linear model with standard errors smoothed by empirical Bayes estimation, taken from the R package limma, to determine differentially enriched proteins. Nominal p-values were transformed using the Benjamini-Hochberg correction to account for multiple hypothesis testing (112). Proteins were considered significantly differentially enriched if they had q values less than 0.01 and an absolute value of log_2_ fold change greater than 1, or twice as enriched linearly.

Data quality were assessed through distance matrices and through principal component analysis. Volcano plots were custom generated but drew heavily from thematic elements from the R package Enhanced Volcano (113). Pathway overrepresentation analysis was performed using MetaScape with default settings (61). Pathway overrepresentation p-values were adjusted using the Benjamini-Hochberg correction and significant pathways were determined from those with q values less than 0.01. Biologically interesting pathways were selected manually, and the gene sets that constituted those pathways were submitted to ProHitz-viz Dotplot generator to view protein-level enrichment differences for the selected pathways (62). In the ProHitz dotplots, the rows were sorted by hierarchical clustering using Canberra distance and Ward’s minimum variance method for clustering. The columns were sorted manually. Venn diagrams for overlapping proteins across the conditions were generated using the R packages ggvenn or ggVennDiagram (114, 115). The heatmap was generated using the R package pheatmap (116). The GO summary table (Table 1) was generated using R package gt (117).

### Immunofluorescence

24 hours post-transfection, cells were washed three times at room temperature with DPBS and fixed in 4% paraformaldehyde for 15 min. After washing, cells were permeabilized in 0.5% Trition-X-100 for 10 min. Cells were then washed three times in either 1x DPBS or 1x Tris-buffered saline (TBS) and blocked in 3% BSA for 1 h at room temperature. After blocking, cells were incubated overnight at 4 °C in primary antibody diluted in blocking buffer. The next day cells were washed three times with DPBS or TBS and incubated in secondary antibody diluted 1:500 or 1:750 in blocking buffer (Cy5 Donkey anti-rabbit, 711-175-152; Cy5 Donkey anti-mouse, 715-175-151; 488 Goat anti-mouse, A-11029). Following incubation, cells were washed three times in DPBS or TBS and mounted onto glass slides using Prolong Gold with DAPI (ThermoFisher; P36935). The following primary antibodies were used: UPF1 (Cell Signaling Technologies; 12040S; 1:2000), MOV10 (Proteintech; 10370-1-AP; 1:1000), VPS35 (Cell Signaling Technologies; 81453S; 1:500), eIF2α (Cell Signaling Technologies; 9722S; 1:500), G3BP1 (Proteintech; 13057-2-AP; 1:2500), PABP1 (Cell Signaling Technologies; 4992S; 1:500), CLTA (Proteintech; 10852-1-AP; 1:500), Twin-Strep-tag® (IBA Lifesciences; 2-1517-001; 1:1000); and Streptavidin 660 Conjugate (ThermoFisher Scientific; S21377; 1:500). Images were collected on a Leica DMi8 THUNDER Inverted Fluorescence Microscope with a DFC7000 T camera (Leica).

### Immunoprecipitation

24 hours post-transfection, cells were washed two times on ice with DPBS. Cells were lysed on ice in either a low salt HEPES buffer (10 mM HEPES, 50 mM NaCl, 5 mM EDTA) or Pierce IP lysis buffer (25 mM Tris/HCl pH 7.4, 150 mM NaCl, 1 mM EDTA, 1% NP-40, 5% glycerol) with 1% Halt protein/phosphatase inhibitor (ThermoFisher Scientific; 78446). Samples were spun at 17,100xg for 15 minutes at 4°C. Protein concentration were measured in the detergent soluble protein fraction by BCA assay (Pierce). Cell lysate was immunoprecipitated with Magstrep Type3 beads (IBA Lifesciences; 2-4090-002) overnight with end/end rocking at 4°C following the protocol provided by the manufacturer. Bound material was eluted from beads in Buffer BXT (0.1 M Tris/HCL pH 8.0, 0.15 M NaCl, 1 mM EDTA, 0.05M Biotin) + β-Mercaptoethanol (BME) at 95°C for 5 minutes. 10% Input, eluted material, and flow-through was then subjected to SDS/PAGE and Western blotting as described below.

### Western Blot

Cell lysis and western blotting was performed as previously described with minor modifications (49). In brief, cells were lysed on ice in either RIPA Buffer (50 mM tris pH = 8.0, 150 mM NaCl, 0.1% SDS, 1% trition-x-100, 0.5% sodium deoxycholate) or cytoplasmic lysis buffer (50 mM tris pH = 8.0, 150 mM NaCl, 0.5% tri-tion-x-100) with 1% protein/phosphatase inhibitor (ThermoFisher; 78442). The RIPA lysate was sonicated and centrifuged for 15 min at 14,000 rpm at 4 °C. The cytoplasmic lysate was vortexed and centrifuged for 15 min at 14,000 rpm at 4 °C. The supernatant was saved as the detergent soluble protein fraction. Protein concentration were measured in the detergent soluble protein fraction by BCA assay (Pierce). Next, cell lysates were analyzed for relative protein expression using SDS/PAGE followed by two-channel infrared quantitative western blots as described previously (47). The samples were denatured in 1X Laemmli loading buffer with 5% tris(2-carboxyethyl) phosphine (TCEP) at 70°C for 15 min. Equal amounts of protein were loaded into a 4–20% PROTEAN TGX Precast Gels (Bio-Rad). After transferring to 0.2 μm nitrocellulose membranes, some blots were stained with Revert 700 (LI-COR; 926–11,010) to measure total protein for normalization and signal was captured at 700 nm on an Odyssey Fc Imaging System (LI-COR), and then destained following the manufacture’s protocol. Protein blots were then blocked in EveryBlot Blocking Buffer (Bio-Rad; 12010020) for 5 minutes at room temperature and incubated with primary antibodies (diluted in blocking buffer) overnight at 4 °C. Membranes were washed three times for five minutes in TBST and then incubated with the appropriate secondary antibody diluted in blocking buffer for 60 min at room temperature. Lastly, membranes were washed three times with TBST for five minutes and visualized using the Odyssey Fc Imaging System (LI-COR). The following primary antibodies were used: StrepMAB-Immo (anti-Twin-Strep-tag®; IBA Lifesciences; 2-1517-001; 1:4000), FUS (1:2000; Bethyl Laboratories; A300-302A), UPF1 (Cell Signaling Technologies; 12040S; 1:1000), MOV10 (Proteintech; 10370-1-AP; 1:800), VPS35 (Cell Signaling Technologies; 81453S; 1:1000), eIF2α (Cell Signaling Technologies; 9722S; 1:500), G3BP1 (Proteintech; 13057-2-AP; 1:2000), PABP1 (Cell Signaling Technologies; 4992S; 1:1000), CLTA (Proteintech; 10852-1-AP; 1:1000), G3BP1 (Proteintech; 13057-2-AP; 1:2000), TAF-15 (Bethyl Laboratories; A300-308A); EWS (Epitomics; 3319-1; 1:1000), Anti-Puromycin (Sigma-Aldrich; MABE343; 1:5000), LC3A/B (Cell Signaling Technologies; 12741; 1:1000); SQSTM1/p62 (Cell Signaling Technologies; 5114; 1:1000), GAPDH (Cell Signaling Technologies; 2118; 1:10,000), and H3 (Millipore; 06-599; 1:5000).

### Quantitative PCR (qPCR)

48 hours post transfection, cells were harvested for RNA using TRIzol™ Reagent (ThermoFisher Scientific; 15596026) following manufacturer guidelines. Equal amounts of RNA were used to create the cDNA library using the High-Capacity cDNA Reverse Transcription Kit with RNase Inhibitor (ThermoFisher Scientific; 4374966). qPCR was performed on a CFX96 Touch Real-Time PCR Detection System (Bio-Rad) using the PowerUp™ SYBR™ Green Master Mix (ThermoFisher; A25741). Results were quantified using the ΔΔCT method. Primers are listed in Supplemental Table 2.

### SUnSet Assay

Puromycin was obtained from Gibco suspended in 20 mM HEPES pH 6.7. Drug was aliquoted and stored at −20°C (ThermoFisher Scientific; A1113803). 48 hours post transfection, cells were treated with 1 µM puromycin diluted in cell culture media for 30 minutes at 37°C/5% CO_2_. Control cells were treated with vehicle (20 mM HEPES pH 6.7) diluted in cell culture media for 30 minutes at 37°C/5%CO_2_. Following treatment, cells were lysed in RIPA lysis buffer+1% protein/phosphatase inhibitor and subjected to SDS/PAGE and Western blotting as described above.

### Autophagosome assay

Bafilomycin A1 (Baf) was obtained from Tocris (#1334) and resuspended in dimethyl sulfoxide (DMSO) and aliquoted and stored at −20°C. 48 hours post-transfection, cells were treated with 0.1 µM Baf diluted in cell culture media for 4 hours at 37°C/5% CO_2_. Control cells were treated with vehicle (DMSO) diluted in cell culture media for 4 hours at 37°C/5%CO_2_. Following treatment, cells were lysed in RIPA lysis buffer+1% protein/phosphatase inhibitor and subjected to SDS/PAGE and Western blotting as described above.

### Statistical analysis

Non-proteomic statistical analysis was performed using GraphPad Prism 8 (San Diego, CA). Effect of variant on FUS localization was determined using an ordinary one-way analysis of variance (ANOVA) with Tukey’s post-hoc test (Figures 1C/D). Effect of variant on UPF1 mRNA fold change expression was determined using an ordinary one-way ANOVA with Tukey’s post-hoc test (Figure 5A). Effect of variant on mRNA fold change for other targets was determined using a two-way ANOVA with Tukey’s post-hoc test (Figure 5B). Effect of variant on autophagosome markers was determined using a mixed model two-way ANOVA with Tukey’s post-hoc test (Figures 5F/G). Significance was reached at p < 0.05. Significance is designated as p < 0.05 (*), p ≤ 0.0021 (**), p ≤ 0.0002 (***), p ≤ 0.0001 (****). All quantified blots either normalized to total protein (Figures 1E, 5D/F/G), GAPDH (Figures 1D/E), or H3 (Figures 1D/E).

## Data availability

The APEX2 mass spectrometry proteomic data from this publication have been deposited to the ProteomeXchange Consortium via the PRIDE partner repository (https://www.ebi.ac.uk/pride/archive/) and assigned the dataset identifier PXD026578 (118).

## Acknowledgements

We thank all the members of the Kukar lab and the Emory Center for Neurodegenerative Disease (CND) for their support and helpful comments during this research. This work was supported by the National Institutes of Health (NIH)/NINDS grants (R01 NS093362, R01 NS105971), a New Vision Research Investigator Award, the Alzheimer’s Drug Discovery Foundation (ADDF), and the Association for Frontotemporal Degeneration (AFTD), the Bluefield Project to Cure Frontotemporal Dementia, and the BrightFocus Foundation to Thomas Kukar.

## Author contributions

*Michelle A. Johnson:* Conceptualization, Methodology, Investigation, Writing-Original Draft, Visualization. *Thomas A. Nuckols:* Methodology, Data analysis/Bioinformatics, Visualization. *Paola Merino:* Methodology, Investigation, Visualization. *Pritha Bagchi:* Methodology, Investigation, Data analysis/Bioinformatics. *Srijita Nandy*: Data analysis/Bioinformatics, Visualization. *Jessica Root:* Methodology, Investigation. *Georgia Taylor:* Methodology, Investigation. *Nicholas T. Seyfried:* Resources, Writing-Review & Editing. *Thomas Kukar:* Conceptualization, Methodology, Funding acquisition, Writing-Review & Editing, Supervision.

## Supporting Information

Supplemental Figures 1-5.

