## Supplemental Figures+Legends for "Proximity-based labeling reveals DNA damage-induced N-terminal phosphorylation of fused in sarcoma (FUS) leads to distinct changes in the FUS protein interactome"

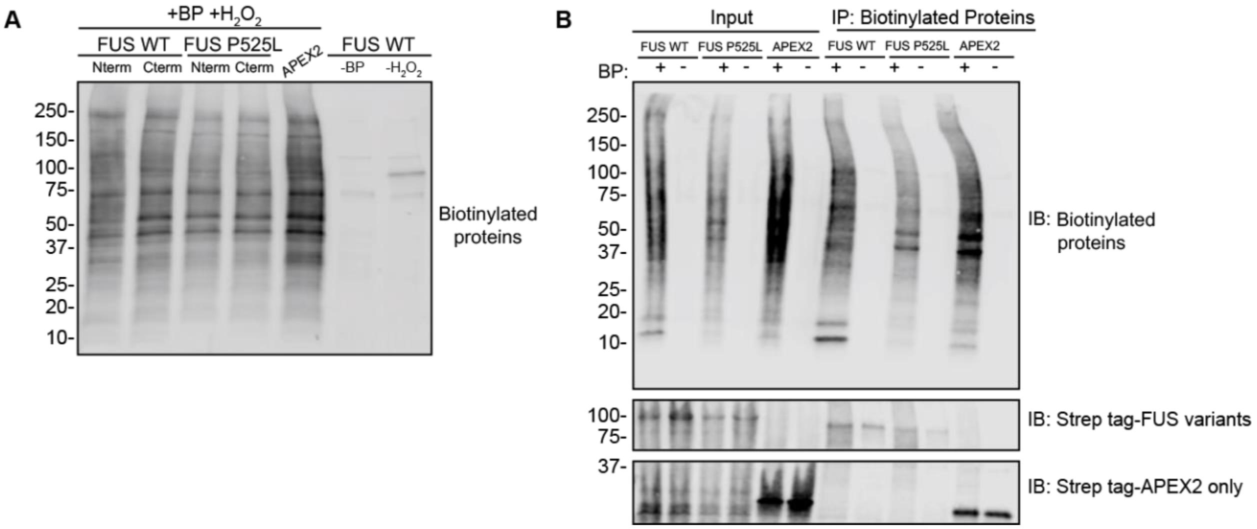


**Supplemental Figure 1 Cells expressing APEX2-FUS fusion constructs must be given biotin-phenol (BP) and H2O2 to induce biotinylation**

*(A) Western blot of cell lysate from HEK293T cells expressing different APEX2-FUS fusion constructs. Constructs either had APEX2 fused to the N-terminus of the FUS variant (Nterm) or the C-terminus of the FUS variant (Cterm) or did not have FUS fused to APEX2 (APEX2). Following transfection, cells were 1) given biotin-phenol (BP) and $H_2O_2$ (+BP + $H_2O_2$), given only $H_2O_2$ (-BP) or given only BP (-$H_2O_2$). 24 hours post-transfection, biotinylation was induced, quenched and cells lysate was harvested and analyzed for biotinylated proteins (streptavidin). (B) Immunoprecipitation of biotinylated proteins using magnetic beads coat in streptavidin showing biotinylated proteins can only be pulled down when cells were given biotin-phenol (BP). Input is 10% of sample loaded onto magnetic beads coated with streptavidin; Elute is 100% of sample eluted off beads. Samples are from cells expressing the following APEX2-FUS fusion proteins: wildtype FUS (FUS WT), P525L FUS (FUS P525L), and APEX2 without FUS fusion (APEX2). Input and elution were analyzed for biotinylated proteins (streptavidin) and Twin-Strep-tag® (strep tag).*

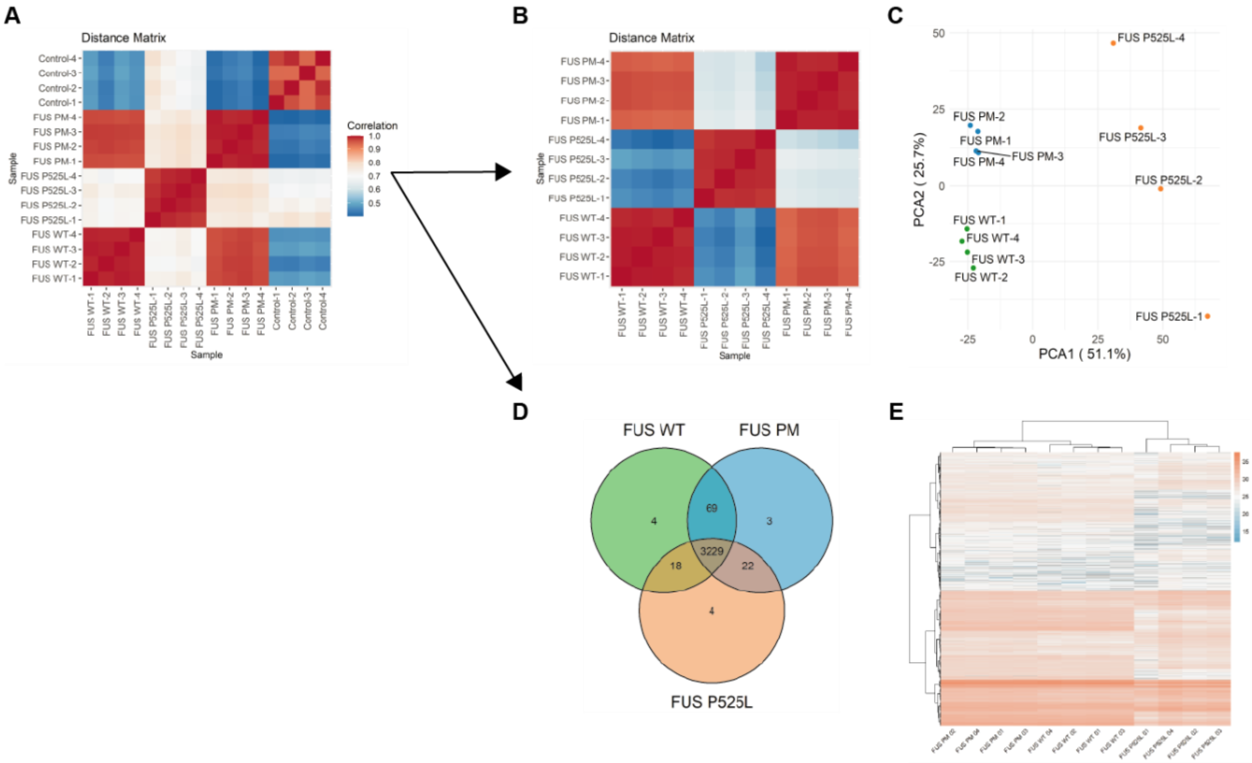


**Supplemental Figure 2 Clustering of protein hits reveals specificity between FUS variant groups**

*(A) Distance matrix of all samples, including controls, showing the Pearson correlation between samples within and between groups with red indicating values closer to 1.0 and blue indicating values closer to 0.5. (B) Distance matrix of all samples, following normalization to control samples, showing the Pearson correlation between samples within and between groups with red indicating values closer to 1.0 and blue indicating values closer to 0.5. (C) Principal Component Analysis (PCA), excluding controls, showing reproducibility of data between biological replicates in three FUS variant groups. (D) Venn diagram of overlap of all protein hits for the three FUS variant groups. (E) Hierarchical clustering of samples based on the intensity profiles of all proteins identified. Missing values are colored gray.*

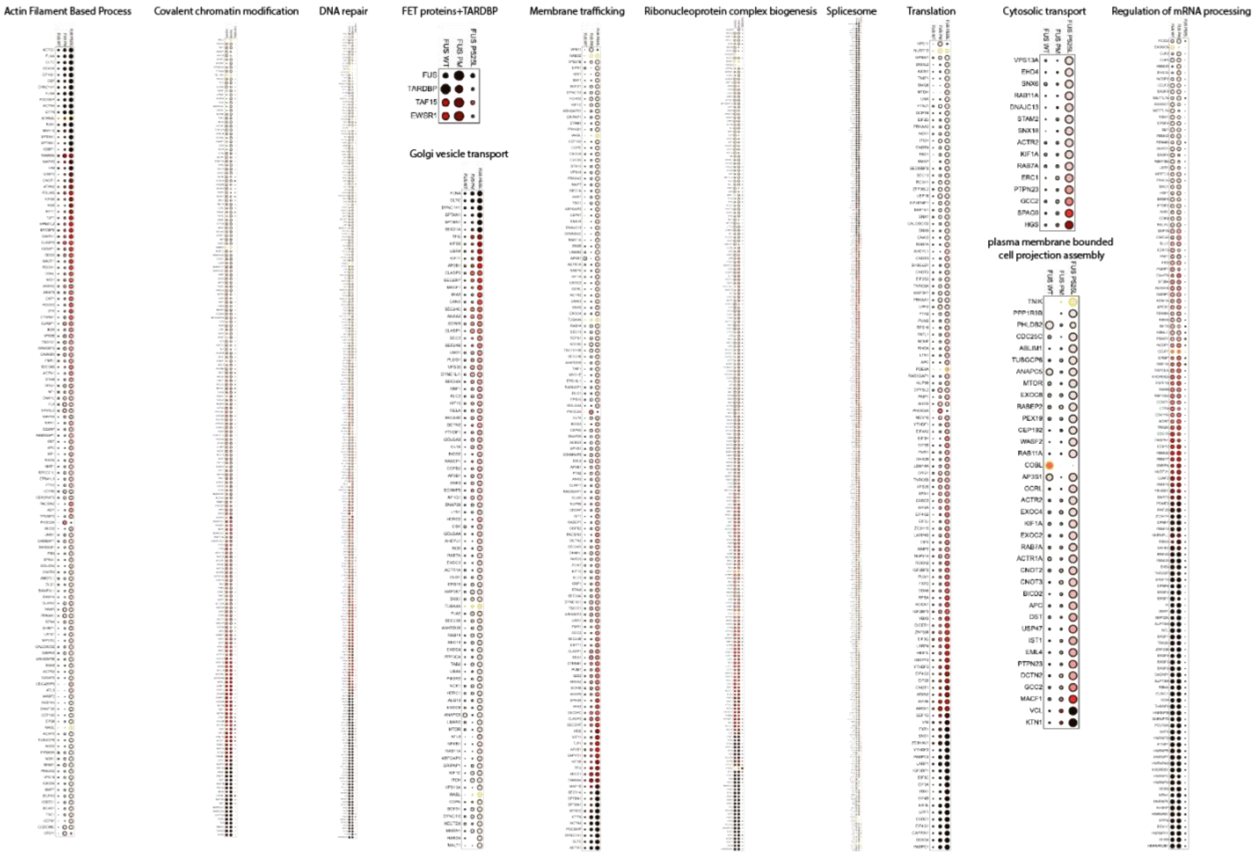


**Supplemental Figure 3 Relative intensity of proteins hits that are relevant for identified ontologies.** *Dot plots were generated using Prohits-viz are a graphical representation of the relative binding intensity of each protein against the three FUS variant groups.*

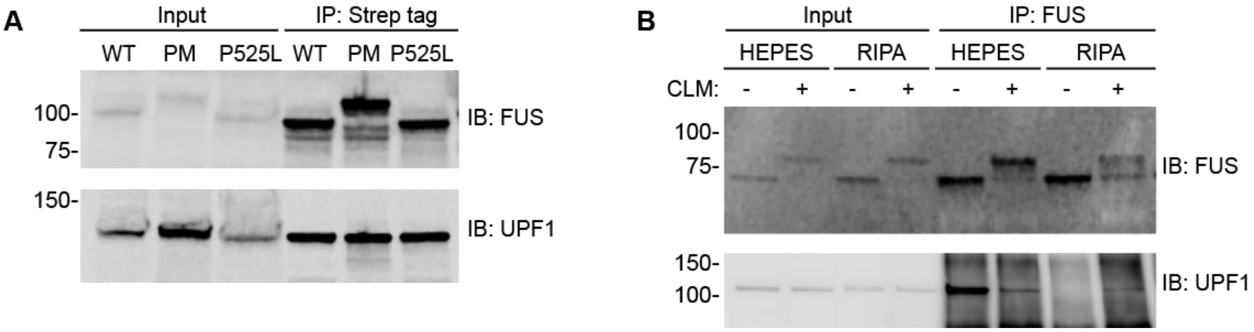


**Supplemental Figure 4 Representative immunoprecipitation of FUS variants.**

*(A) Representative western blot (IB) showing clear enrichment of FUS variants following immunoprecipitation of Twin-Strep-tag® compared to input lysate. Input lysate is 10% of sample loaded Magstrep Type3 beads. (B) Immunoprecipitation for FUS in cells treated with either calicheamicin γ1 (CLM) (+) or vehicle (DMSO; (-)). HEK293T cells were treated with 40 nM of CLM for 3 hours at 37˚C/5% CO2. Cells were lysed either in HEPES (120 mM NaCl, 40 mM HEPES pH 7.4, 0.3% (w/v) CHAPS) or RIPA (150 mM NaCl, 1% NP-40, 0.5% sodium deoxycholate, 0.1% SDS, 50 mM Tris/HCl pH 8.0) based lysis buffer + protein/phosphatase inhibitor. Following lysis, equal amounts of protein were loaded onto Protein G Dynabeads (Thermofisher Scientific, 10003D) coated in FUS antibody (Bethyl Laboratories; A300-302A) for overnight capture and bound material was eluted off following manufacturer instructions. Input and elution were immunoblotted for listed targets.*

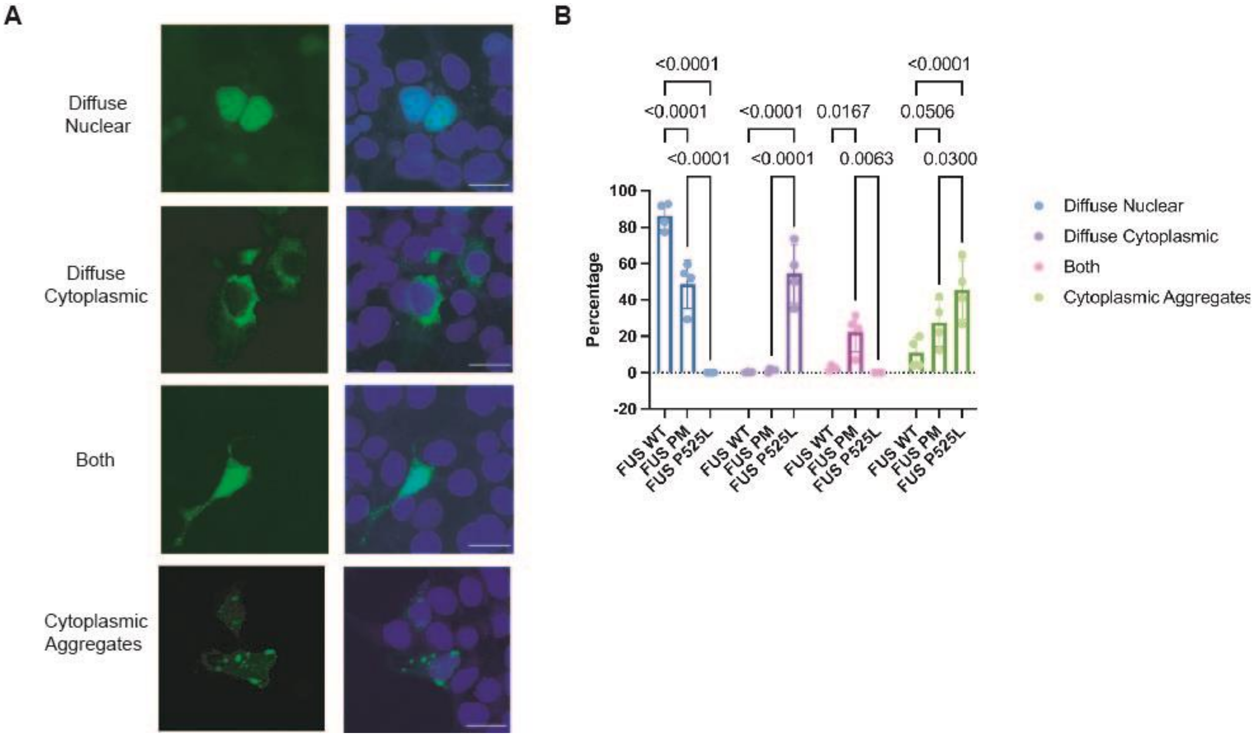


**Supplemental Figure 5 FUS PM forms more cytoplasmic aggregates than FUS WT.**

*Cells were transfected with a GFP-tagged FUS variant (FUS WT, FUS PM, and FUS P525L). (These are the variants used in Figures 4 and 5) On average 165 cells were then classified into one of four categories based on the localization of FUS: Diffuse Nuclear (signal was spread diffusely throughout the nucleus), Diffuse Cytoplasmic (signal was spread diffusely throughout the cytoplasm), Both (signal was spread diffusely throughout the nucleus and cytoplasm) and Cytoplasmic Aggregates (signal was present in cytoplasmic punctate). The boundaries of nucleus were determined using the DAPI signal. (A) Representative images of the four classifications schemes for FUS localization. (B) The percentage of cells in each category was calculated for each group and a two-way ANOVA was performed to determined significance (n=4). Error bars indicate mean ± SEM.*
